# Randomizing the human genome by engineering recombination between repeat elements

**DOI:** 10.1101/2024.01.22.576745

**Authors:** Jonas Koeppel, Raphael Ferreira, Thomas Vanderstichele, Lisa M. Riedmayr, Elin Madli Peets, Gareth Girling, Juliane Weller, Fabio Giuseppe Liberante, Tom Ellis, George M. Church, Leopold Parts

## Abstract

While protein-coding genes are characterized increasingly well, 99% of the human genome is non-coding and poorly understood. This gap is due to a lack of tools for engineering variants that affect sequence to the necessary extent. To bridge this gap, we have developed a toolbox to create deletions, inversions, translocations, and extrachromosomal circular DNA at scale by highly multiplexed insertion of recombinase recognition sites into repetitive sequences with CRISPR prime editing. Using this strategy, we derived stable human cell lines with several thousand clonal insertions, the highest number of novel sequences inserted into single human genomes. Subsequent recombinase induction generated an average of more than one hundred megabase-sized rearrangements per cell, and thousands across the whole population. The ability to detect rearrangements as they are generated and to track their abundance over time allowed us to measure the selection pressures acting on different types of structural changes. We observed a consolidation towards shorter variants that preferentially delete growth-inhibiting genes and a depletion of translocations. We isolated and characterized 21 clones with multiple recombinase-induced rearrangements. These included viable haploid clones with deletions that span hundreds of kilobases as well as triploid HEK293T clones with aneuploidies and fold back chromosomes. We mapped the impact of these genetic changes on gene expression to decipher how structural variants affect gene regulation. The genome scrambling strategy developed here makes it possible to delete megabases of sequence, move sequences between and within chromosomes, and implant regulatory elements into new contexts which will shed light on the genome organization principles of humans and other species.

## Introduction

The human genome is a vast desert of non-coding sequence sprinkled with small islands of DNA that contain the instructions to make proteins. These protein-coding sequences only make up ∼1% of our genomes and are interrupted by an average of 8 introns that need to be actively removed upon transcription (*1*). Another ∼8% of our genome consists of non-coding sequences with biochemical marks correlating with regulatory activity (*2*) and the majority (54%) of the genome is repetitive (*3*). It remains unclear how much DNA in our genomes is dispensable for cellular survival and how the expression of genes changes when manipulating the order and position of nearby sequences.

The importance of genome structure, order, and content can be probed by comparing how alternative genome configurations behave in cells. Experimentally, site-specific recombinases have the capacity to induce diverse DNA sequence alterations such as deletions, inversions, and translocations which have shed light on genomic organization principles (*4–6*). However, the application of recombinases has been primarily confined to investigating individual loci and not the entire human genome. Alternatively, complex genome shattering events can be observed in pathologies such as chromothripsis in cancer, and illuminate the space of viable genome configurations (*7*, *8*). These drastic alterations can lead to an unexpected increase in cellular adaptability including drug resistance (*9*). Despite the instructive nature of these natural genomic phenomena, our observations are confined to post-selection results and we remain without the means to initiate these rearrangements at will or monitor them in real-time.

A method to generate variants at genome scale and in an experimentally controllable way has been pioneered in baker’s yeast, *S. cerevisiae*, through the incorporation of hundreds of symmetrical loxP sites (loxPsym) into synthetic chromosomes. Each of the sites could act as an anchor for recombination by Cre recombinase. Collectively, these sites form the substrate of an inducible genome evolution system called SCRaMbLE (synthetic chromosome rearrangement and modification by loxP-mediated evolution) (*10–12*). This approach is uniquely targeted, avoids double-strand breaks, and preserves exon integrity. A wealth of knowledge about genome rearrangements has been obtained through this system. For example, scrambling one synthetic chromosome and moving genes across varying expression neighborhoods affected isoform selection (*13*) while scrambling six synthetic chromosomes linked rearrangement patterns with chromatin accessibility and 3D arrangements, connecting chromatin structure with genome evolutionary dynamics (*11*). However, the human genome is vastly larger compared to the yeast genome, putting the synthesis of chromosomes for SCRaMbLE currently out of reach. In addition, the sparsity of genes in the human genome would likely result in a much different landscape of tolerated variation.

Scrambling human genomes would require inserting recombinase sites into the genome at scale. Prime editing is a novel genome-editing technique that makes it possible to precisely engineer short sequence insertions without double-strand DNA breaks and external DNA donor templates (*14–16*). To achieve the necessary scale, recurring patterns of repetitive elements offer an opportunity to insert many recombinase sites simultaneously (*17*). For example, long-interspersed elements-1 (LINE-1) are a class of transposable elements that make up 17% of the human genome. Previous studies have targeted these abundant repetitive elements with base editors and demonstrated the feasibility of highly multiplexed genome editing (*18–20*).

Here, we combine the SCRaMbLE and prime editing technologies to generate human genomes with vastly changed structure, content, and ploidy (Figure 1A). We report stable cell lines with thousands of loxPsym insertions into LINE-1 retrotransposons, despite the considerable toxicity associated with manipulating numerous loci across different cell lines. We characterize these prime-edited cell lines and explore the features of LINE-1s that make them amenable to editing. Scrambling cell lines with hundreds of integrated recombinase sites generates thousands of diverse structural variants. By comparing variants generated shortly after induction and those remaining in culture after two weeks, we map the selection pressures acting on these variants. In the process we discover megabase scale deletions that can survive in haploid genomes and characterize changes in gene expression in cell clones with multiple Cre-induced rearrangements. Our findings make substantial progress in elucidating the selective forces shaping structural changes of the genome and provide a strategy to manipulate mammalian genomes at previously unseen scale.

**Figure 1.**
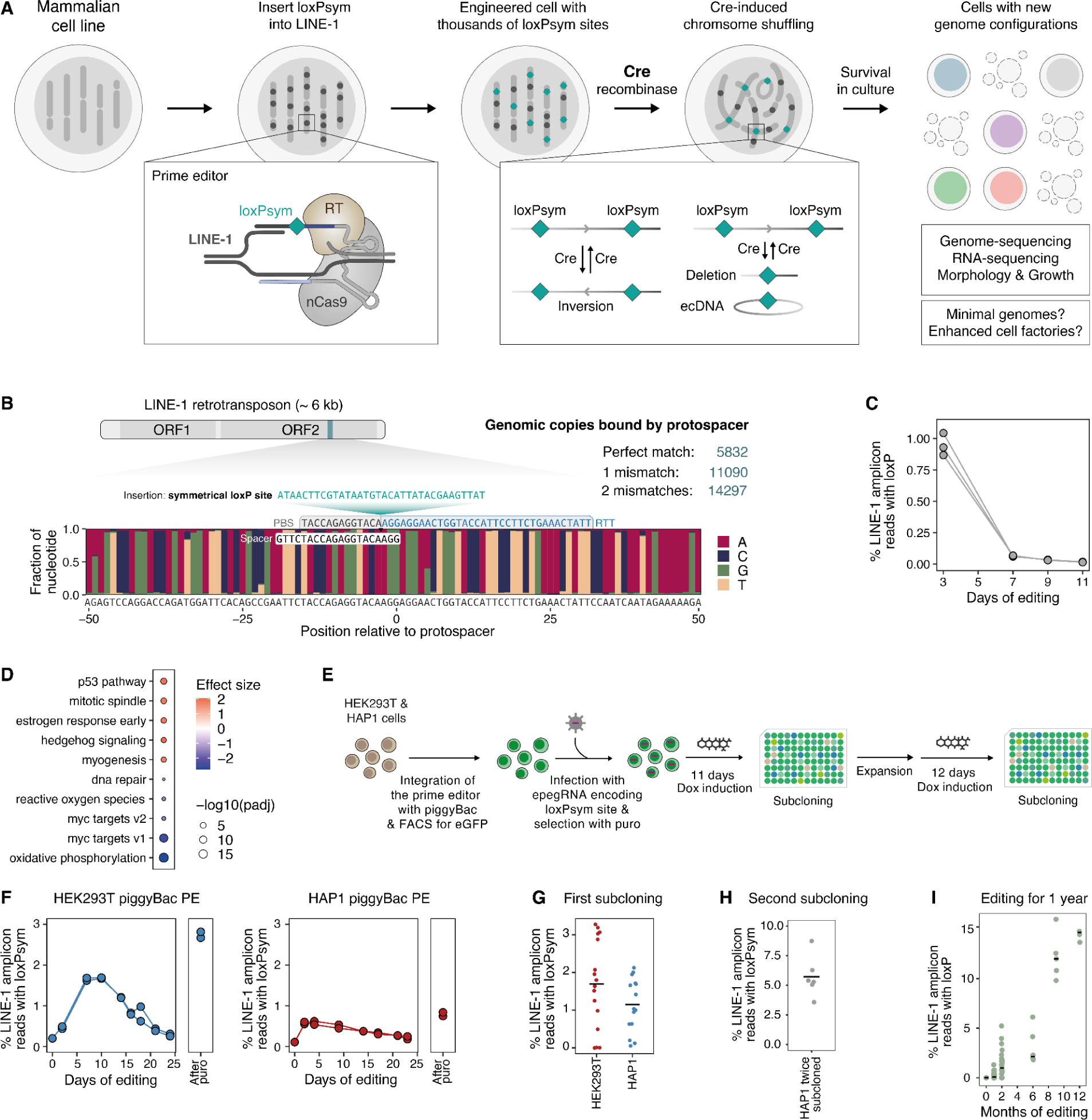
Highly multiplexed prime editing to insert hundreds of recombinase sites. (**A**) Schematic of a strategy to scramble mammalian cells. Thousands of loxPsym sites are inserted into LINE-1 with prime editing. Induction of Cre recombinase shuffles the chromosomes. Surviving derivative clones are sequenced and phenotyped. (**B**) Schematic of a LINE-1 retrotransposon with two open reading frames (ORF), the target site, and protospacer of the pegRNA in blue. Lower panel: Nucleotide frequencies (y-axis) in a 100 bp window around the nicking site (x-axis) for all 14,297 LINE-1 sequences with 2 or fewer mismatches to the protospacer. The positions and sequences of the primer binding site (PBS) and reverse transcriptase template (RTT) including loxPsym insertion and homology arm are indicated. (**C**) Frequencies of LINE-1 amplicon reads with loxPsym insertions (y-axis) over 11 days (x-axis) following transfection of the prime editing reagents. Markers and lines show one of N = 3 biological replicates. (**D**) Top and bottom 5 differentially expressed pathways (by p-value) between cells transfected with prime editor alone or with prime editor and the LINE-1 targeting pegRNA. N = 2 biological replicates. (**E**) Schematic of an editing time course with two subcloning steps to enrich cells with hundreds of loxPsym insertions. (**F**) Frequencies of LINE-1 amplicon reads with loxPsym insertions (y-axis) over 25 days of editing and after selection with puromycin at the end of the time course (x-axis) in HAP1 and HEK293T cells with stably integrated prime editor and engineered pegRNAs (panels). Markers and lines show one of N = 2 biological replicates. (**G**) Frequencies of LINE-1 amplicon reads with loxPsym insertions (y-axis) for HEK293T or HAP1 clones (markers) after one round of subcloning. (**H**) Frequencies of LINE-1 amplicon reads with loxPsym insertions (y-axis) for HAP1 clones (markers) after two rounds of subcloning. (**I**) Frequencies of LINE-1 amplicon reads with loxP insertions (y-axis) for HEK293T clones (markers) that were sequentially grown out and subcloned over the course of 12 months (x-axis).

## Results

### Stable cell lines with hundreds of recombinase site insertions

We hypothesized that prime editors could simultaneously insert hundreds of recombinase sites into the genome of a cell if targeted to high copy-number elements. The feasibility of large-scale genome editing at repeating sequences in mammalian cell lines has been demonstrated by Smith et al., who targeted base editors to LINE-1s introducing more than 10,000 mutations in a single genome (*18*). Their best-performing gRNA targeted a highly conserved region in the second open reading frame (ORF2) of LINE-1 (Figure 1B) and had 5,832 perfect matches to the human reference genome (17,496 in a triploid genome). We converted this gRNA into a prime editing gRNA (pegRNA) by adding a 3’ extension containing a reverse transcriptase template with 34 nt overlap to the LINE-1 sequence, a loxPsym site, and a 13 nt primer binding site (Figure 1B) and co-transfected the pegRNA with prime editor 2 (PE2) into HEK293T cells. We observed successful editing of 1.5% of all targeted LINE-1s (average of 200-300 insertions in triploid genomes of the population) after two days. However, the fraction of edited elements plummeted to only 0.02% by day 11 (Figure 1C), indicating that prime editing of a high copy number element was detrimental to survival.

The nicking Cas9 in prime editors creates single-strand DNA breaks and thousands of simultaneous nicks could trigger the DNA damage response in the edited cells. Indeed, RNA sequencing (RNA-seq) of cells transfected with prime editor and the LINE-1 targeting pegRNA identified the upregulation of pathways associated with the DNA damage response, such as the p53 pathway (Figure 1D). Concurrently, there was downregulation in growth-related pathways, including Myc signaling and the oxidative phosphorylation pathway suggesting that cells might be slowing down growth in response to DNA damage. Similarly, Smith et al. observed toxicity when editing LINE-1s with nicking, but not with nuclease-dead base editors. Using nuclease-dead Cas9 is not currently possible for prime editing due to the chemistry of the reaction, which requires a single-strand break for priming.

While a burst of high prime editor activity from transient transfection overwhelms the cell, lower levels of editing and accumulation of loxPsym insertions into LINE-1s over a prolonged period of time may be tolerated. To enable this, we integrated multiple copies of doxycycline-inducible prime editors into HEK293T and near-haploid, fibroblast-like HAP1 cell lines (*21*) using the piggyBac transposon system (*22*) (Supplementary Figure 1A). These prime editing lines can insert a symmetrical loxP site into a single locus (HEK3) with up to 80% efficiency after 10 days (Supplementary Figure 1B). We further improved efficiency by using engineered pegRNAs (epegRNAs) that introduce a protective RNA pseudoknot motif to avoid 3’-end degradation and adding the p53-inhibitor piftherin-alpha and basic fibroblast growth factor (bFGF) (*23*, *24*).

Putting these ideas together, we infected the stable, prime editing HEK293T and HAP1 cell lines with lentivirus encoding the LINE-1-targeting epegRNA, selected for lentiviral integration with puromycin, and induced editing with 1 µM doxycycline and 10 µM piftherin-alpha (Figure 1E). We monitored the insertion rates every 3-4 days and observed an increase in edited LINE-1s over the first 2-5 days, peaking at an average of 1.7% on day 10 for HEK293T cells and 0.62% on day 4 for HAP1 cells followed by a gradual decline (Figure 1F). The apparent decline could be caused by cells that inactivated the prime editor or the epegRNA and outcompete the cells that are still capable of editing. Indeed, selection for active expression from the epegRNA cassette using puromycin in the cell population after 32 days killed more than 90% of all cells (Supplementary Figure 1C) and increased the average fraction of edited LINE-1s to 2.7% in HEK293T cells and 0.79% in HAP1 cells (Figure 1F) corresponding to the insertion of hundreds of recombinase sites.

We reasoned that single cell sorting after a period of active editing could separate actively editing cells from cells which silenced the prime editing machinery (Figure 1E). After 11 days of editing, we derived and analyzed 16 clones for HEK293T cells and 15 clones for HAP1 cells, and observed editing rates between 0 and 3.3% for HEK293T cells and 0 and 2.1% for HAP1 cells (Figure 1G). To further increase the number of loxPsym site insertions, we subjected one HAP1 clone with 1.9% initial editing to another 12 days of doxycycline induction followed by a second round of subcloning which resulted in up to 8% LINE-1 editing (Figure 1H).

To probe the limits to adding insertions to LINE-1, we grew a HEK293T cell line continuously for one year and subcloned it every 3-4 months. We always picked the subclone with the highest editing rates and kept expanding it. Ultimately, we derived a clone that had edited 16% of all targetable LINE-1s (Figure 1I). This cell line contains the highest number of single sequence insertions into the human genome that we are aware of and demonstrates that it is possible to continuously engineer genomes at massive scale using prime editing. RNA-seq of individual clones taken at 0, 6, 10, and 12 months during our long-term editing strategy revealed that Myc targets, oxidative phosphorylation, fatty acid metabolism, and apoptosis are the most significantly regulated pathways, corroborating our initial findings (Supplementary Figure 2, Figure 1D).

To assess the feasibility of implementing this strategy in non-cancerous diploid cells, induced pluripotent stem cells PGP1 (Personal Genome Project: GM23338) and human skin fibroblasts were subjected to the same engineering process. As with HAP1 and HEK293T, we sequentially integrated the dox-inducible prime editor followed by the epegRNA with puromycin and blasticidin resistance, respectively (Supplementary Figure 3A). Following a 17-day period of doxycycline induction, the resulting surviving cells underwent single-cell cloning. From the PGP1 population, we derived three unique clones, while the fibroblasts yielded 22 distinct clones. For both cell types, sequencing revealed only minimal insertion of loxP sites, with insertion rates reaching a maximum of 0.028% for PGP1 and 0.0036% for the fibroblast population (Supplementary Figure 3B), corresponding to 7 and 1 insertions on average.

### Characterization of edited cells with hundreds of loxP insertions

To precisely map the locations and number of integrated loxPsym sites, as well as unintended genome alterations, we whole-genome sequenced three edited clones (referred to as HAP1-loxPsym(301), HEK293T-loxPsym(638), and HEK293T-loxPsym(1697)) using long-read technology (Oxford Nanopore, Figure 2A). We mapped 301 clonal insertions in the HAP1 and 638 and 1697 sites in the two HEK293T clones. Besides loxPsym, pegRNA, prime editor, and other insertions, we detected no translocations, and only 7 additional structural variants in the HAP1-loxPsym(301) clone and 30 in the HEK293T-loxPsym(638) clone that are not already present in parental HAP1 and HEK293T cells (Supplementary Figure 4A). In addition, the read depth profile of edited HAP1s were consistent with a single allele of all chromosomes except for a segment of chr15, in line with the parental genotype (*21*) (Supplementary Figure 4B). The lack of variants and copy number changes demonstrated that month-long editing and insertion of hundreds of loxPsym sites did not cause large-scale genome instability in the whole-genome sequenced clones.

**Figure 2:**
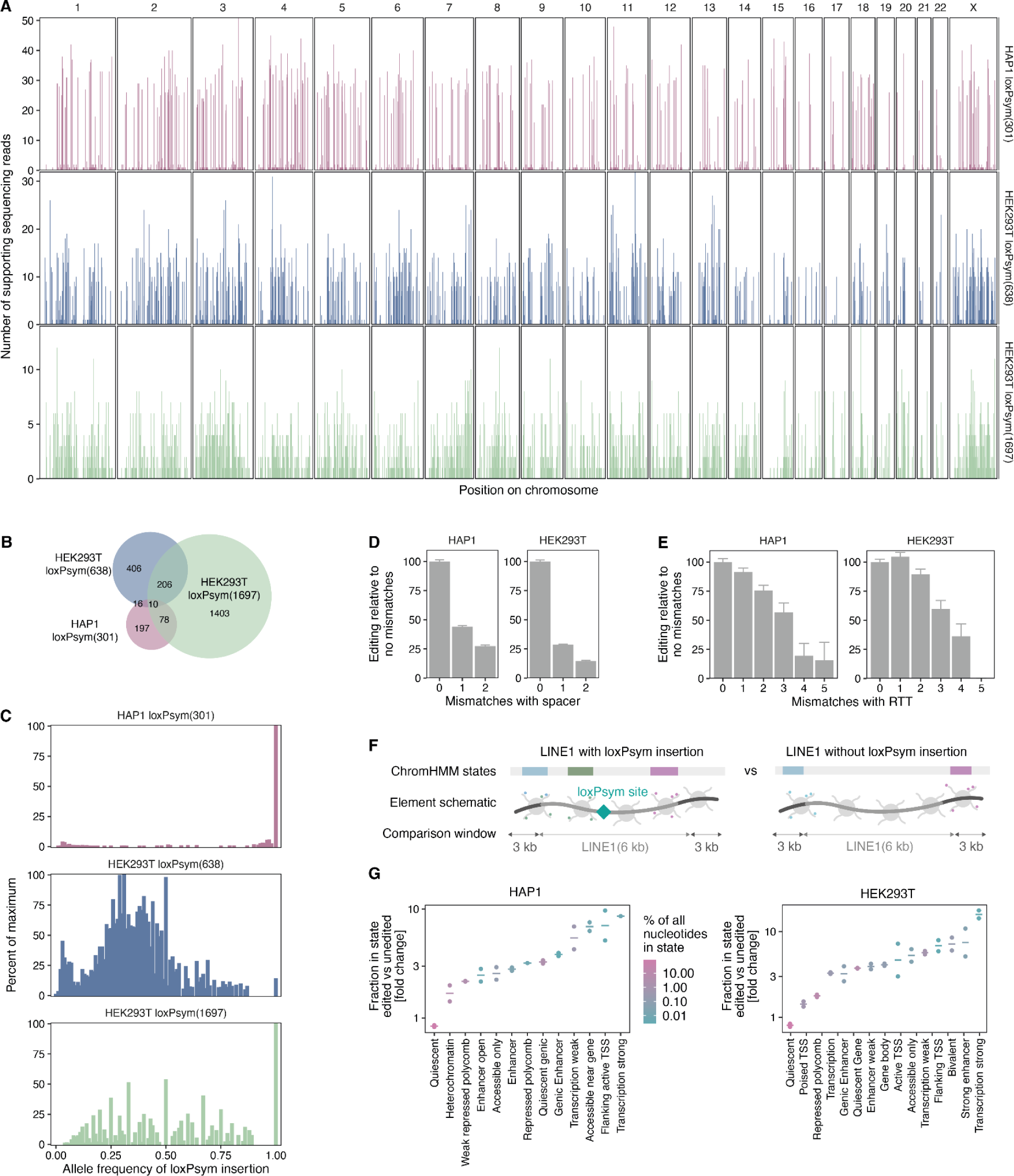
Cell lines with hundreds of loxPsym sites distributed across their genomes. (**A**) Locations of loxPsym site integrations (vertical bars) and the number of supporting sequencing reads (y-axis) across chromosome positions (x-axis) for three clones (panels). (**B**) Overlap of loxPsym sites between the three sequenced clones (colors, circles). The area of the Venn diagram is proportional to the number of loxPsym sites. (**C**) The abundance of loxPsym insertions (percent of the allele frequency with maximum abundance, y-axis) at various allele frequencies (x-axis) in the three sequenced clones (panels, colors). (**D**) Average sequencing reads from targeted sequencing of edited LINE-1s normalized to no mismatches (y-axis) for LINE-1s with different numbers of mismatches to the pegRNA protospacer (x-axis) across two cell types (panels). Error bars: standard error of mean. N = 2 biological replicates. (**E**) As (D) but for LINE-1s with mismatches with the reverse transcriptase template (x-axis). Sum of N = 2 biological replicates. (**F**) Schematic of the comparison in chromatin composition between edited (n > 1 read) and unedited (N = 0 reads) LINE-1s. (**G**) Fraction of edited to non-edited LINE-1s (y-axis) for overall nucleotides in chromatin state (x-axis) in HAP1 and HEK293T cells (panels) colored by the fraction of all nucleotides in the human genome that are in a given state. Cross bars represent averages from N = 2 biological replicates (markers).

The insertions were broadly distribution across the genome with 14.3 sites on average per chromosome (range 1-34) for the HAP1 cells and a strong correlation with the number of LINE-1s with perfect matches to the pegRNA (HAP1: R^2^=0.83, HEK293T: R^2^=0.90, Supplementary Figure 4C). Insertions across the three clones mapped to largely different LINE-1s and only 10 were shared across all (Figure 2B). The median allele frequencies of loxPsym site insertions were 1.00 for HAP1-loxPsym(301) cells and 0.35 for HEK293T-loxPsym(638) cells, consistent with their haploid and mostly triploid genomes, respectively. The HEK293T-loxPsym(1697) cells had a median allele frequency of 0.5 indicating occurrence of multiallelic integrations (Figure 2C).

LINE-1 sequences represent sources of near-identical sequences that could be used to understand variation in prime editing efficiencies. To map insertion sites at higher throughput, we devised a Cas9-enrichment-based long-read sequencing method that targets the Cas9 nuclease to LINE-1s with loxPsym insertions (*25*, *26*) (Methods) (Supplementary Figure 5). Elements with up to two mismatches to the protospacer were still successfully edited at 25 days, but at lower frequencies (27% and 15% in HAP1 and HEK293T cells, Figure 2D). In contrast, one and two mismatches in the 34 nt homology arm of the reverse transcriptase template were well tolerated, but editing rates dropped for elements with three or more mismatches (Figure 2E).

Finally, to evaluate the impact of chromatin on prime editing, we trained the ChromHMM model on eight chromatin tracks (Supplementary Figure 6A. Methods) and correlated its predicted abundance of chromatin states in the LINE-1s and 3 kb of surrounding sequence to the editing rate (Figure 2F). Editing events were most enriched in LINE-1s with transcription and enhancer-related chromatin signatures, and depleted in quiescent (no chromatin marks) or repressed/heterochromatic elements (Figure 2G). Some chromatin states only make up small fractions of the genome and clustering of LINE-1s based on all chromatin marks showed the same trend; edited LINE-1s were enriched in clusters associated with transcription and depleted of ones associated with quiescent intergenic DNA (Supplementary Figure 6I,J). This result was also robust to changes in the window sizes of the surrounding sequence considered (Supplementary Figure 6i,j) and is consistent with recent work exploring chromatin context for prime editing (*27*, *28*).

### Scrambling the human genome

To scramble the genome of our clones with hundreds of loxPsym sites, we treated the cells with membrane-permeable TAT-Cre protein for 4 hours, recovered for 16 hours, and used long-read sequencing to map the Cre-induced rearrangements (Figure 3A, Supplementary Figure 7, Methods). At this early time point, selection pressures would not have had time to deplete variants that are detrimental to survival or chromosome segregation. Each read with a rearrangement likely represents an independent recombination, since the complexity of the cell pool is much higher than the sequencing coverage, the cells had no time to replicate, and the DNA was not amplified for sequencing. The number of rearrangements scales linearly with the number of loxPsym sites per chromosome, both for variants within the same chromosome (*cis*) (R^2^=0.92/0.95 in HAP1 and HEK293T) and across different ones (*trans*) (R^2^=0.83/0.77, Supplementary Figure 8). With 50x and 23x genome coverage, we detected 6,481 and 1,166 rearrangements in HAP1 and HEK293T cells, respectively (Figure 3B). Assuming haploidy in HAP1 and triploidy in HEK293T, Cre treatment induced 131 or 143 rearrangements per cell on average.

**Figure 3:**
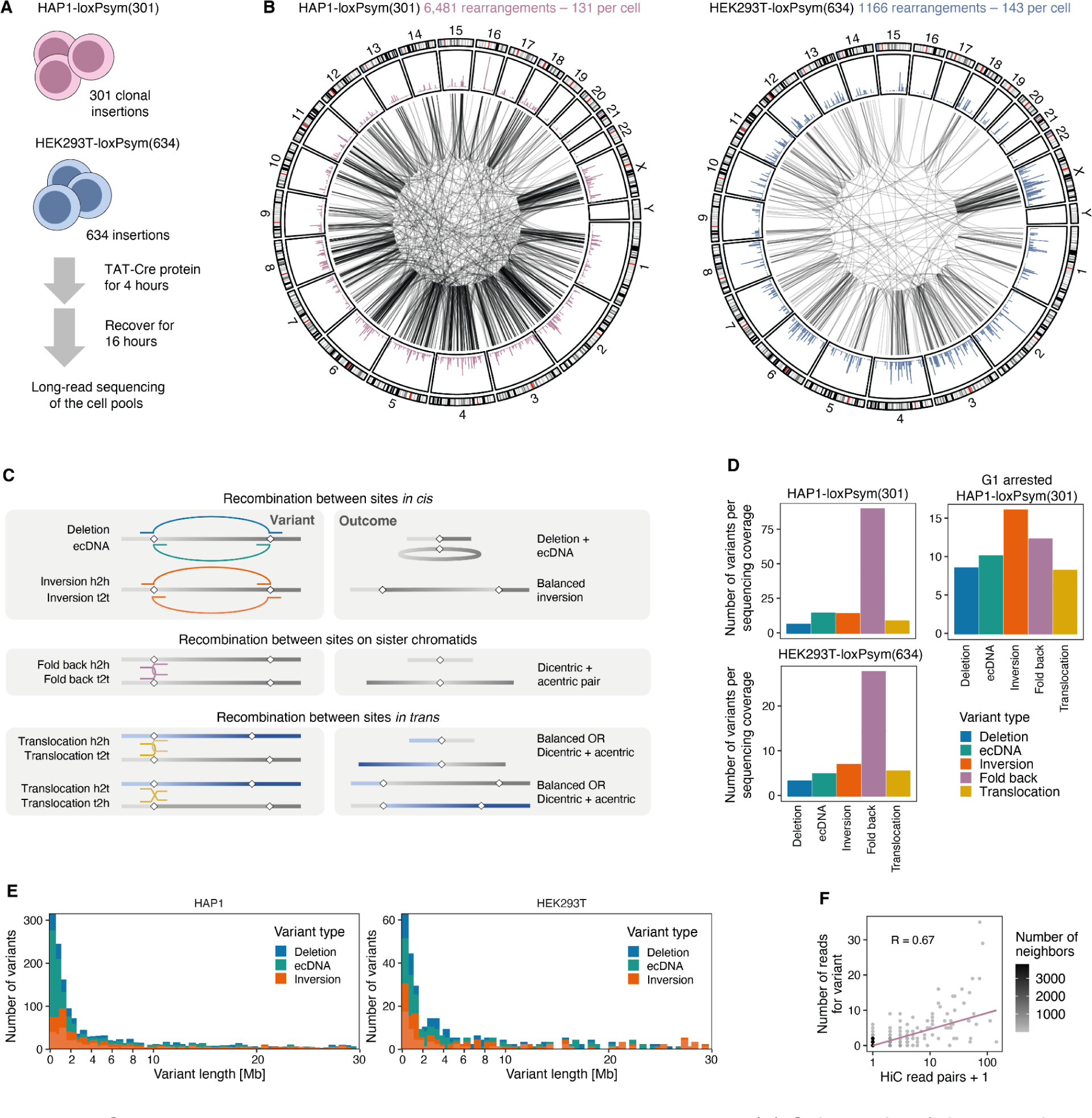
Cre recombinase induced thousands of rearrangements. (**A**) Schematic of the experiment. (**B**) Overview of all structural variants produced in the two clones after 4 hours of scrambling and 16 hours of recovery. From outside to inside: Chromosomes (ideograms), integration locations and rearrangement frequencies for loxPsym sites (bars), Cre induced structural variants (arcs). (**C**) Schematic of the classes of variants that can be generated by Cre induction. Gray and blue lines represent chromosome sequence according to the reference genome with diamonds indicating the locations of loxPsym sites. Arcs represent rearrangements that join two DNA segments of varying orientation and location. Color according to variant class. h2h: head to head, t2t tail to head. (**D**) The types of structural variants produced (x-axis, color according to C) and their counts (y-axis) and separated by cell line and whether they were inhibited in G1/S during Cre induction (panels). (**E**) The number of structural variants in *cis* (y-axis) stratified by the distance between the two breakpoints (x-axis). (**F**) Number of supporting sequencing reads (y-axis) for cis-variants in HAP1 cells (markers) with different number of read pairs in an independent Hi-C experiment in HAP1 cells (x-axis). Line: linear regression fit.

Whenever Cre recognizes two loxPsym sites and catalyzes a rearrangement, the outcome can fall into different classes based on the orientation of the DNA ends joined and whether this occurs in *cis* or in *trans*. We can distinguish these different events based on the mapping of sequencing reads at breakpoints (Figure 3C). If two loxPsym sites are located in *cis*, Cre can produce a pair of inversions (10% or 14% of rearrangements in HAP1 and in HEK293T cells respectively, Figure 3D), or a pair consisting of a extrachromosomal circular DNA (ecDNA, HAP1: 11%; HEK293T: 10%) and a deletion (HAP1: 4.5%; HEK293T: 6.6%). In contrast, if the two sites are located on two non-homologous chromosomes in *trans*, the outcome will be a translocation (HAP1: 10%; HEK293T: 11%).

Curiously, the most common class of variants that we observed were fold backs (HAP1: 68%; HEK293T: 58%), a type of rearrangement that creates a U-turn of sequence at the loxPsym site resulting in a ‘mirrored’ inversion (Figure 3C). Fold backs could arise between homologous chromosomes or two sister chromatids of the same chromosome after replication (Figure 3C). The fraction of fold backs was indeed reduced 3.1-fold, down to 22% of total variants, when HAP1-loxPsym(301) cells were arrested in G1 using the CDK4/6 inhibitor palbociclib (Methods, Supplementary Figure 9) prior to Cre induction (Figure 3D). Therefore, cell cycle modulation can alter the types of variants generated. This is important since fold backs that generate a pair of acentric (no centromere) and dicentric (two centromeres) chromosomes are catastrophic for cell survival due to chromosome segregation errors during mitosis. Overall, the frequency of variant types was similar between the two cell lines despite their different ploidy (Figure 3D).

The frequency of deletions and inversions decreased with increasing distance between loxPsym sites, with 71% or 63% of variants shorter than 10 Mb for HAP1 and HEK293T cells (Figure 3E). The median variant length was 3.0 or 4.9 Mb (HAP1/HEK293T), which in HAP1 cells is 15.7 times smaller than the expectation of 47 Mb assuming equal probability of recombination for any given pair of loxPsym sites on the same chromosome (Methods, Supplementary Figure 10A). The observed data could be predicted better by reducing the expected frequency of two loxPsym sites rearranging by 7.5% for each Mb distance between them (exponential decay model, Supplementary Figure 10B,C), consistent with previous observations that the efficiency of Cre-induced rearrangements decays with distance (*6*).

The exponential decay model of generating rearrangements in *cis* predicts a dearth of long variants; however, despite this we do observe a tail of very long deletions and duplications (Figure 3E). This suggests that other factors, such as 3D proximity, may influence the probability of observing a rearrangement. A model that uses 3D distance, assessed by counting Hi-C read pairs, fits the data better (Pearson’s R = 0.68 (Hi-C, (*29*)) vs 0.55 (length), Figure 3F). A linear model with both linear distance and Hi-C pairs only modestly outperformed one with only Hi-C pairs (R = 0.68 vs 0.67) suggesting that 3D contact frequency between two recognition sites is a better explanation of Cre efficiency. Overall, these results are consistent with a recent study by Zhou et al. profiling 260,000 Cre-induced rearrangements in SCRaMbLEd yeasts (*11*) which observed that rearrangement frequency was driven by contact frequency and DNA accessibility.

### Selection pressures shape surviving variants

To understand how selection acts on the various types of rearrangements generated after Cre induction, we grew out the HAP1 and HEK293T cells that survived the scrambling process for 13-15 days and long read sequenced the cell pools at 53-164 times whole genome coverage. In addition, we single-cell sorted cells that showed evidence of recombination based on an integrated reporter, and sequenced 21 expanded colonies. We observed a 188-fold (HAP1) or 87-fold (HEK293T) depletion of variants in the pools over two weeks of growth (Figure 4A), implying strong selection against the majority of rearrangements. After selection, an average of 0.7 rearrangements remained per HAP1 cell (assuming haploidy) or 1.6 per HEK293T cell (assuming triploidy). Similarly, cells that rearranged an integrated Cre activity reporter (and switch from dsRed to tBFP expression upon recombination, Figure 4B) depleted from 68% at day 1 to 1% at day 10 suggesting strong selection against cells that took up Cre (Figure 4C). The depletion was less pronounced for both the reporter and remaining variants in the pool (2.4 on average per cell) if the cells were arrested in the G1 phase with palbociclib before Cre induction (Figure 4A,C), consistent with the generation of fewer fold backs (Figure 3E).

**Figure 4:**
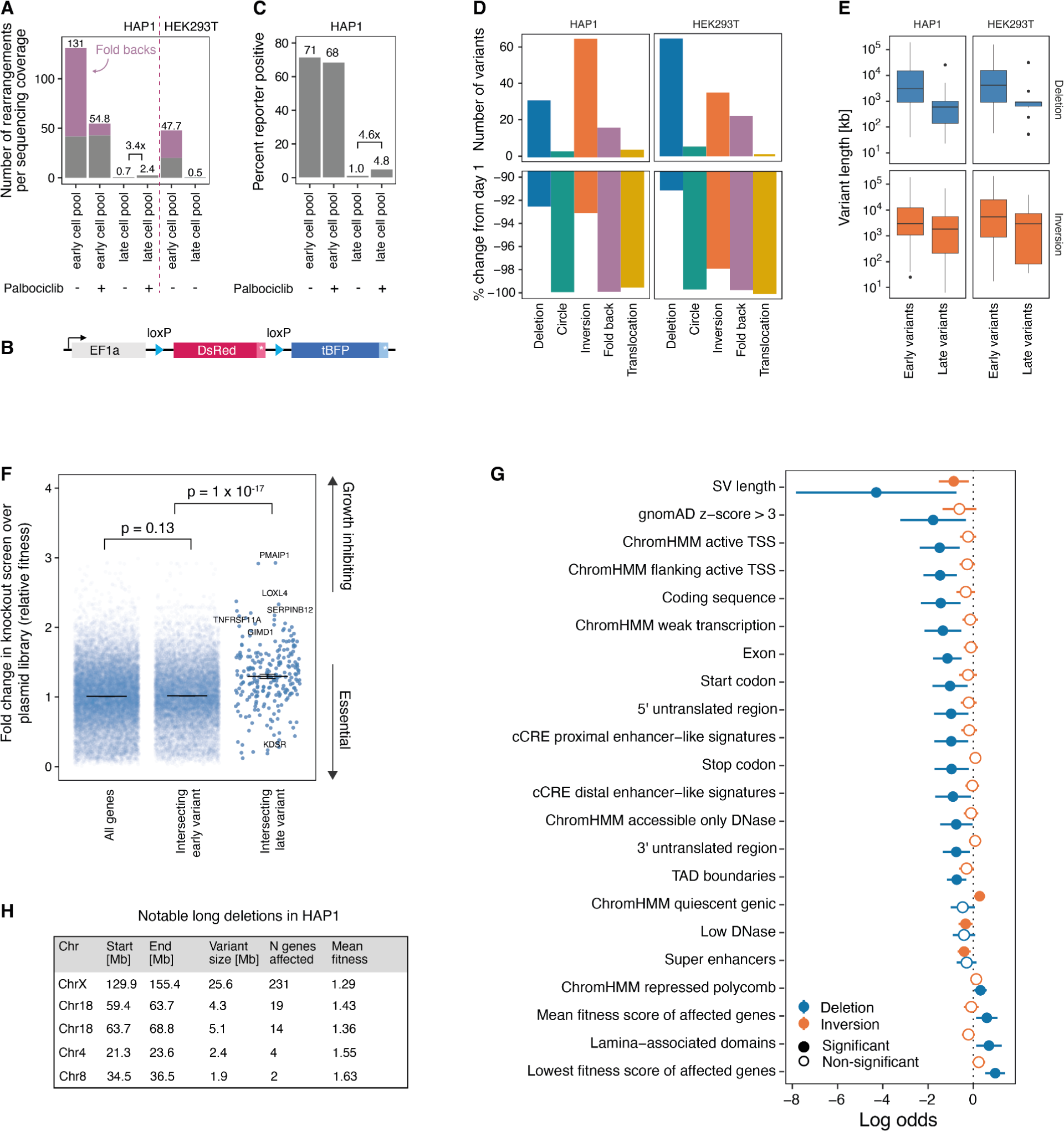
Surviving variants are shorter, depleted of translocations and preferentially delete growth-inhibiting genes. (**A**) Number of total Cre-induced rearrangements per sequencing coverage (y-axis) for early and late cell pools that were untreated or inhibited in G1 prior to scrambling (x-axis). Bars are colored by variant type (lighter blue: fold backs, darker blue: all other types). (**B**) Schematic of the Cre reporter. (**C**) Percent of cells that were BFP-positive in flow cytometry (y-axis) after rearranging a genome-integrated Cre reporter (schematic bottom) at different time points (x-axis). The cells were either restricted in G1/S with palbociclib or not prior to scrambling (x-axis). (**D**) Upper panel: The types of structural variants produced (x-axis) and their counts (y-axis) in HAP1 and HEK293T cells (panels) at 13-15 days after Cre induction. Lower panel: The percent difference in variant abundance per sequencing coverage compared to day 1. (**E**) Variant length (y-axis) for early and late variants (x-axis) stratified by cell line and variant type (panels). Box: median and quartiles. Whiskers: Largest or smallest value no further than 1.5 interquartile ranges from the hinge. (**F**) Average fold changes of four guides targeting a gene (markers) in a HAP1 CRISPR knockout screen (y-axis) for all genes, genes intersecting early deletions (day 1) or intersecting late (day 13-14, x-axis). (**G**) Log odds (x-axis) for features (y-axis) between early variants and surviving variants colored by variant type. Markers represent the log odd and whiskers 95% confidence intervals. Unless indicated otherwise, the fraction of each variant covered by the respective feature is used as the statistic. Only significant features shown. (**H**) An example of five long deletions in HAP1 cells.

The frequency of variant types we observed before (Figure 3D) and after scrambling (Figure 4C) differs starkly. Because Cre joins up all DNA ends of the two loxPsym sites involved in the recombination, all structural variants are initially balanced. However, some rearrangement products lack centromeres (ecDNA and acentric chromosomes from fold backs or translocations), cannot be segregated properly and will be lost during subsequent cell cycles. Indeed, between HAP1 and HEK293T cells, we see a 99.9/99.6% depletion of ecDNAs, a 99.8/99.7% depletion of fold-backs and a 99.5/100% loss of translocations (Figure 4D). However, the depletion of translocations cannot be explained by centromere configuration alone since 50% of translocations should result in two derivative chromosomes with one centromere each. Deletions (HAP1: -93.0%, HEK293T: -97.8%) and inversions (HAP1: -92.4%, HEK293T: -91.0%) were less depleted compared to ecDNAs, fold-backs, or translocations, but fewer than 1 in 10 variants remained.

Deletions might be selected against because they remove essential genes. To test this experimentally, we performed a genome-wide CRISPR knockout screen in wild type HAP1 cells (*30*) (Supplementary Figure 11). We defined relative fitness as the average fold change of gene-targeting guides after two weeks in culture (methods). Relative fitness values <1 indicate a growth-defect and values >1 indicate a growth advantage upon loss. Only 2/17 deletions removed any gene with a >50% growth defect in the HAP1 knockout screen, in contrast to early deletions where 137/185 did (Supplementary Figure 12A). Longer variants have more chance to delete essential DNA and we indeed see a consolidation towards 4.5 to 5.0-fold shorter deletions after weeks in culture for HEK293T and HAP1 cells, respectively (median early 3.0/4.2 Mb vs late 0.60/0.93 Mb, Figure 4E). Moreover, genes contained within persisting deletions were enriched for ones whose loss conferred a growth advantage in the CRISPR knockout screen (average of genes in the deletions = 1.3, t-test p=10^-17^, Figure 4F). A much weaker selection was apparent for inversions (average = 1.08, p = 5x10^-5^, Supplementary Figure 12B).

To better understand the genomic features under selection we compared 822 early and 70 late time point inversions and deletions from HAP1 cells across 49 features spanning chromatin states (Supplementary Figure 6), conservation, mutational constraint, regulatory elements and gene essentiality. Late deletions were significantly depleted in constrained regions (gnomAD z-score), gene-associated features (coding sequences, exons, start and stop codons, untranslated regions), and active regulatory elements (transcription start sites and enhancers) (Figure 4G, Supplementary Figures 13-14). Conversely, late deletions were enriched for genes with higher fitness scores, polycomb repressed regions, and lamina-associated domains, which are usually heterochromatic. As expected, inversions showed weaker signals of selection across all features but were significantly shorter at later time points and depleted of super enhancers. Many of the features are correlated with one another. For example, gene annotations are correlated with active chromatin and anticorrelated with lamina associated domains (Supplementary Figure 13A). To understand if the signal we observed is driven by proximity to essential genes alone, we calculated log odds using a model that contains the relative fitness score of the most essential affected gene in a given variant as an additional variable (Supplementary Figure 13C). While the directionality of features was maintained, all but 7 lost nominal significance. Together this data suggest that the main, but not exclusive, driver of selection against deletions is the loss of essential genes.

Many of the remaining deletions in HAP1 cells were millions of base pairs long (Figure 4H), highlighting that even in haploid cells substantial amounts of DNA are dispensable for growth under cell culture conditions. One 4.3 Mb variant on chromosome 18 deleted 19 expressed (TPM > 0.1) genes with an average relative fitness of 1.43. This variant removed both a selectively essential as well as a strongly growth suppressing gene: 3-ketodihydrosphingosine reductase (KDSR) had a relative fitness score of 0.2 in HAP1 and was selectively essential in the cancer dependency map (183/1095 cell lines (*31*)). Conversely, PMAIP1 is pro-apoptotic gene and its loss was highly growth promoting in the HAP1 screen (relative fitness = 2.9) and in 1035/1095 cancer dependency map cell lines. Curiously, the loxPsym site marking the end for this deletion was the start point for another, independent deletion that covered another 5.1 Mb on chromosome 18 (Figure 4H) but did not contain any essential genes in the CRISPR screen (relative fitness range of 11 deleted genes: 1.0-2.0). Since the sequencing was done from a pool of cells it is possible that some of the variants observed did not happen in isolation or occurred in cells that previously reverted to diploidy (14% of the cell population at the time of sequencing). This is likely the case for one 25.6 Mb variant that deleted 231 genes on chromosome X including many essential ones.

Together, these data demonstrate that continued survival and growth in cell culture consolidated an initially mixed pool of variants towards shorter deletions and inversions. Surviving deletions preferentially affected growth-inhibiting genes and avoided essential genes, regulatory elements, and highly constrained DNA.

### Scrambled clones

Clones that survived the scramble process represent a unique collection of novel human cell lines that only differ in specific structural variants. We sought to leverage this resource to understand how such variants affect gene expression and ultimately cell fitness by generating shallow depth whole genome sequencing for 21 clones and RNA sequencing for three clones with interesting genetic architectures.

The first investigated HAP1 clone had three Cre-induced variants, a 23 kb deletion that did not overlap with any genes, a 599 kb deletion that deleted the *EPHA7* receptor tyrosine kinase gene, and a 40 kb inversion that affected the last three exons of *TMEM38B*, a cation channel involved in calcium homeostasis (Figure 5a, Supplementary Figure 15A,B). *EPHA7* and *TMEM38B* were indeed the top and 21st most downregulated genes (5000 and 24-fold respectively, Figure 5B). For the inversion and deletions, none of the genes within 3 Mb of the structural variant were significantly dysregulated (Figure 5C,D). To measure the effect of the three variants collectively affecting 662 kb of DNA on fitness of the clone, we competed it against parental cells and observed no difference in growth speed (Figure 5E, Clone 1).

**Figure 5.**
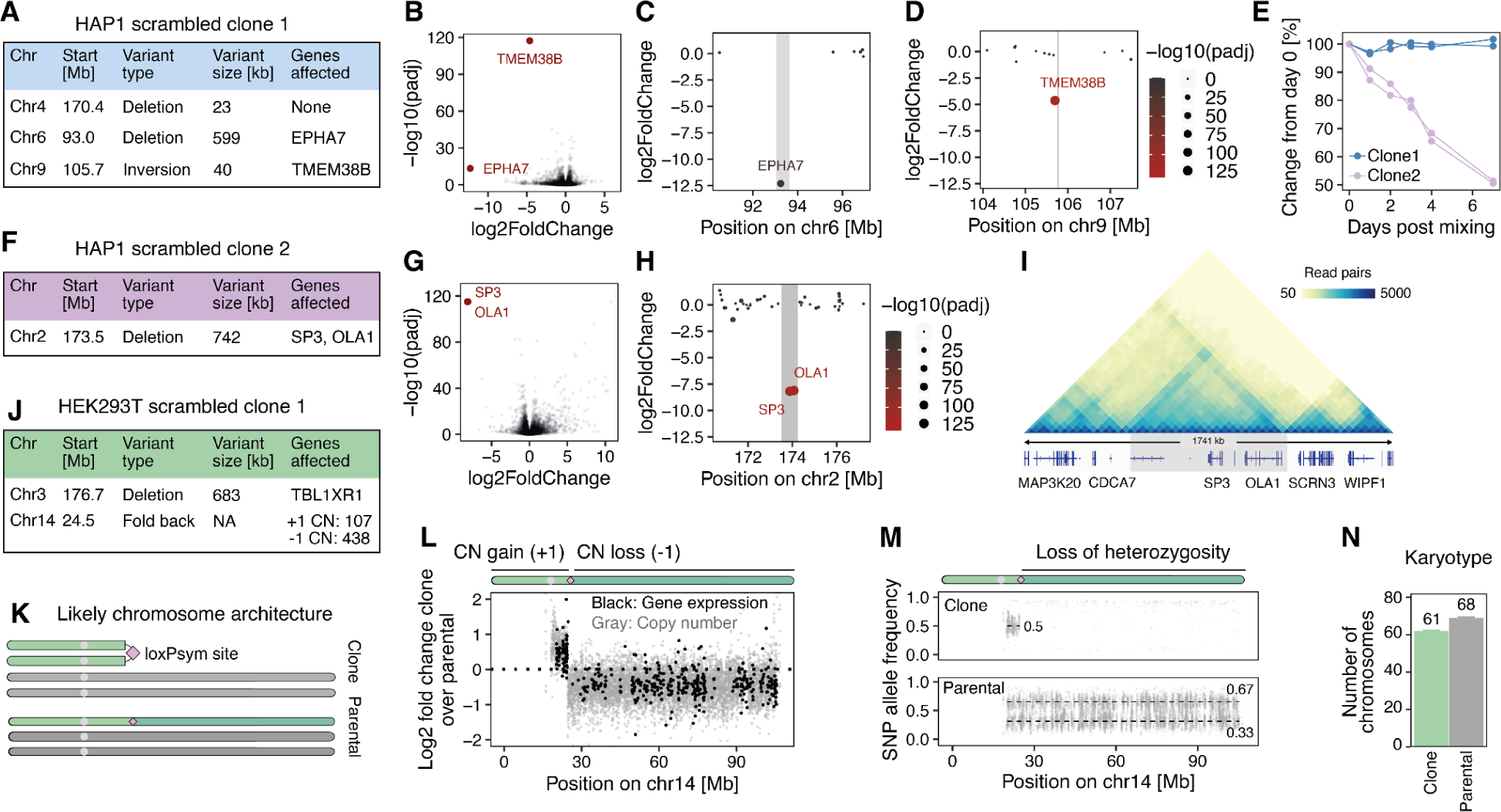
Structural variants in clones directly affect gene expression. (**A**) A table with variants found in the HAP1 scrambled clone 1. (**B**) Significance of differential gene expression (y-axis) and expression fold change (x-axis) between clone 1 and parental HAP1 cells (DEseq2) for genes with at least 10 reads (markers) from N = 2 biological replicates. (**C**) Changes in gene expression (y-axis) between the scrambled clone and parentals for genes (markers) located +/- 3 Mb of a 599 kb deletion (x-axis, deletion shaded gray). Marker size and color according to significance. N = 2 biological replicates. (**D**) As in (C) but around a 40 kb inversion. (**E**) Changes of clone abundance post mixing with parental cells (y-axis) for two scrambled HAP1 clones (colors) with two biological replicates (lines) over 7 days (x-axis). (**F**) Table of variants in HAP1 clone 2. (**G**) As (B) but comparing clone 2 and parentals. (**H**) As (C) but across a 742 deletion in clone 2. (**I**) Triangle heatmap showing the frequency of chromatin interaction read pairs (color gradient) within a 1741 kb segment (deletion shaded gray) and gene annotations below the heatmap. (**J**) A table with variants found in the HEK293T scrambled clone 1. (**K**) Schematic of a fold-back on chromosome 14. The loxPsym site is indicated as a pink diamond, and the centromere as a gray circle. (**L**) Log2-fold changes between clone and parental (y-axis) for mean whole-genome sequencing read depth in 50 kb windows (gray markers) or gene expression (black markers) along chromosome 14 (x-axis). N = 2 biological replicates. (**M**) B allele frequencies (y-axis) for heterozygous SNPs along chromosome 14 (x-axis) for clone and parental (panels). Dashed lines at 0.5 and 0.33. (**N**) Number of chromosomes (y-axis) assessed by karyotyping of 24 metaphase spreads of parental or clone1 cells (x-axis). Bars and numbers indicate average, error bars the s.e.m.

The second HAP1 clone we examined had one 745 kb deletion that affected two genes, *OLA1* and *SP3* which were the top and second most downregulated genes in this cell line (Figure 5F-G). The 10 Mb surrounding the 742 kb deletion are gene-rich and contain 68 expressed genes. However, none of the surrounding genes were significantly misregulated (Figure 5H). Intriguingly, the variant almost exactly excises a single topology associated domain (TAD) containing *OLA1* and *SP3* (Figure 5I). The regulatory elements that were within the TAD (Supplementary Figure 15C) might not be able to influence genes outside the TAD, and consequently, their deletion would not affect neighboring gene expression. *OLA1* is weakly essential (relative fitness: 0.64) and many genes were significantly differentially expressed in this cell line (652 up, 172 down; adjusted p-value < 0.01, Fold change < 0.5 or > 2). Growth promoting pathways (MYC target genes 1 and 2) were downregulated and the cell line had a growth defect compared to parental cells (Figure 5E, Clone 2).

Lastly, one HEK293T scrambled clone had a 683 kb deletion on chromosome 3 and a fold back on the minor allele of the triploid chromosome 14 (Figure 5J,K, Supplementary Figure 16A,B). The fold back should have resulted in a dicentric/acentric pair of isochromosomes. The dicentric derivative chromosome persisted, which raised the copy number for 24.5 Mb of the underlying sequence from three to four (Figure 5L) and changed the single nucleotide polymorphism B allele frequency from 0.33/0.66 to 0.5/0.5 (Figure 5M). The acentric derivative chromosome was lost, resulting in a decrease of copy number for 76.5 Mb of sequence from 3 to 2 (Figure 5L) and loss of heterozygosity (Figure 5M). The genes encoded on the affected sequences followed the expected trend (median increase by 34% for duplicated sequences and decrease by 27% for lost sequences). In addition, the clone lost entire copies of chromosomes 1, 2, 4, 7, 10, 11, and 20 resulting in an average karyotype of 61 chromosomes compared to 68 chromosomes of the parental cell line (Figure 5N, Supplementary Figure 16C-E). These pervasive chromosome losses could have been caused by missegregation after scramble events that created dicentric-acentric pairs. This clone highlights how scramble in cells with higher ploidy can create isochromosomes and aneuploidies, thus forming a useful resource to study how aneuploidy affects growth and gene expression and how dicentric chromosomes are resolved.

## Discussion

In this study, we leveraged prime editing of repetitive elements to engineer the incorporation of thousands of Cre-recombinase sites into the human genome, creating a substrate to induce thousands of distinct recombinations at will. The variants affect all chromosomes and encompass deletions, inversions, ecDNAs, fold backs and translocations. We found that several megabase-scale deletions are viable in haploid cells for growth in culture. More broadly, we defined the characteristics of tolerated structural variation by comparing sequence, gene annotation, regulatory, mutational constraint, and conservation features between generated and surviving variants. Finally, we characterized clones with several large Cre-induced rearrangements, including an isochromosome, and found that variants strongly affected gene expression when they modified copy number but did not influence the expression of nearby genes.

Prime editing is currently the most versatile genome editing technology because it enables direct search and replace operations on DNA. Here, we used it to insert thousands of sequences into the same genome, obtaining engineered cell lines with the highest number of novel sequence insertions that we are aware of. The integrated recombinase sites made it possible to scramble a mammalian genome for the first time, a starting point for forays into the vast space of potential genomes for biotechnology and basic research. Manipulation of genomes at this scale was previously only achievable through *de novo* genome synthesis in a genome 0.3% the size of ours (*32–34*).

Our strategy to scramble the human genome to generate thousands of distinct structural variants for study gives an inroad to illuminate the extensive but understudied non-coding genome. In one such experiment, we detected more than 200 deletions and 500 inversions that together span 4 or 9 gigabases of sequence - several times the size of the entire human genome. So far, most of our knowledge on structural variation comes from heavily pre-selected germline variants in the human population (*35–39*) or somatic variants in cancers (*40*, *41*) that represent the small fraction of all variants that are compatible with survival. In contrast, Cre-induced rearrangements at early time points have not yet entered the selective ratchet, and the vast majority of the initially diverse landscape of genomic alterations did not persist over the course of two weeks in culture. Still, the remaining deletions and inversions together affected over 3% and 12% of the genome, respectively – more than the combined total of all protein-coding sequences. The advantage of our strategy is that the persisting variants can be directly compared against the baseline of all generated ones to illuminate the selection pressures acting on the genome. For example, we saw that the remaining variants were enriched in quiescent and heterochromatic regions and avoided coding regions as well as gene regulatory elements.

Alongside deletions, inversions, translocations, and fold backs, Cre recombination also created hundreds of ecDNAs. Most of these ecDNAs lack centromeres and cannot be actively separated upon cell division. Accordingly, they did not persist in the context of our transformed human cell lines. Yet, ecDNAs are common and associated with poorer prognosis in human cancers where their uneven distribution across daughter cells can boost copy numbers into the hundreds, amplifying underlying oncogenes such as *EGFR*, *MDM2*, and *MYC* (*42*, *43*). A recent study used a similar Cre-lox system to engineer ecDNAs in primary cells and demonstrated that ecDNAs containing *MDM2* would accumulate over the course of several weeks and help the primary cells overcome p53-dependent senescence (*44*). Genome-wide scrambling in primary cell types could identify regions that are positively selected for when amplified as ecDNA, and shed light on the genesis, propagation and biology of ecDNAs.

We engineered, scrambled, and characterized the effects of structural variants and aneuploidies on the survival and gene expression for two cell lines. While insights from these lines will be highly valuable for bioproduction (HEK293T) and basic research (HAP1), both are transformed cells with abnormal karyotypes. Regulatory sequences are highly context-specific (*45*) and sequences that can be deleted in one cell line might be essential in another context. To generalize towards more contexts, we attempted to expand our method to stem cells (fibroblasts and iPSCs). While transformed cells were able to tolerate highly multiplexed prime editing, non-transformed cells exhibited substantial cytotoxicity. This suggests that non-transformed cells may possess distinctive DNA repair mechanisms or heightened stress responses upon extensive nicking, consistent with our previous work showing that catalytically inactive Cas9 is required for successful editing in iPSCs using the same gRNA (*18*). Notably, HAP1 and HEK293T cells harbor multiple mutations known to inhibit apoptosis and promote cell survival (*21*, *46*) and future endeavors in stem and primary cells would require strategies to prevent apoptosis in response to extensive genomic nicking. Nevertheless, the HAP1 model system has been instrumental for the classification of pathogenicity in protein variants (*47–49*) and our findings on the properties of tolerated structural changes should also generalize more broadly.

The size, number and diversity of variants that we can create is a function of the number of integrated loxPsym sequences. In the HAP1-loxPsym(301) clone, recombinase sites are separated by a median of 7 Mb, expected to contain over 40 genes, several of which are essential. This is in contrast to the synthetic yeast chromosomes where loxPsym sites are only separated by a few hundred base pairs (*10*). The distance of recombinase sites will decrease with more inserted sites as is the case for the HEK293T cell that we cultured for 1 year where the median distance was 2.5 Mb. In addition, several improvements to prime editing and pegRNA design have been published in the meantime (*27*, *50–52*) which should all make it easier to insert a higher number of sequences with our strategy. While our investigation focused on LINE-1s, other repetitive elements (Alu sequences, endogenous retrovirus, and various types of microsatellites) could also be targeted (*19*, *23*). An approach incorporating multiple types of repetitive elements, each with a unique genomic distribution, could enhance the number of rearrangement anchors and achieve a more diverse landscape of rearrangements.

Unlike evolution through point mutations, structural changes generated by scrambling can simultaneously affect tens to hundreds of genes and megabases of non-coding regions. By exploring a much larger mutational space genome scrambling enables a more comprehensive assessment of a phenotypic landscape. We envision that these properties open up exciting possibilities in two major applications.

Firstly, scrambling can generate novel cell lines with evolved properties. By coupling scrambling as a source of sequence diversity to phenotypic selection we can optimize cellular properties and analyze the resulting evolutionary paths to better understand genotype-phenotype landscapes. These types of experiments have already been successful in yeast strains with synthetic chromosomes (*53–56*). In mammalian cells, scrambling could shed light on mechanisms of drug-resistance, growth under adverse conditions (e.g. in minimal medium), or help in biomanufacturing. For example, the HEK293 derivative cell lines are used to produce the ChAdOx1 nCoV-19 vaccine (*57*), as well as a wide range of proteins (*58*).

Secondly, the random generation of large deletions opens up the exciting possibility of assaying the essentiality of sequences genome-wide. With an increasing diversity and density of recombinase site positions across a population of cells, the generation of 10,000s of deletions spanning the genome many times over becomes feasible. By measuring variant dropout, we can build genome-wide maps of essentiality. Such essentiality maps, like the maps being generated from saturation genome editing studies (*59*), will be invaluable for the interpretation of pathogenic variants and provide a deeper understanding of genome structure and function.

## Supporting information

Insertion sites of loxPsym and loxP sequences in one HAP1 and two HEK293T clones

Cre-induced variants at in HAP1 and HEK293T cells at early and late time points and in scrambled clones

Insertion frequencies of loxPsym sequences in LINE-1s with up to 2 mismatches

ChromHMM states in HAP1 and HEK293T cells

Read count tables for RNA sequencing experiments performed in this study

Essentiality of genes in the HAP1 cell assessed by knockout screening

## Acknowledgements

E.M.P., J.K., J.W., F.G.L., G.G., T.E, and L.P. were supported by Wellcome (grant no. 220540/Z/20/A, ‘Wellcome Sanger Institute Quinquennial Review 2021–2026’). R.F. was supported by the Knut and Alice Wallenberg Foundation (KAW 2019.0581). G.M.C, L.M.R, and R.F were supported by the Aging and Longevity-Related Research Fund and EGL Charitable Foundation. T.V. was supported by a BBSRC iCASE Studentship (BB/V509425/1). We thank the Sanger flow cytometry core facility, sequencing pipelines, and sequencing walkup teams for supporting us in the study. We thank the BPF Genomics and the HMS Immunology Flow Cytometry Core Facilities at Harvard Medical School for their expertise and instrument availability.

## Author contributions

Conceptualized and initiated the study: J.K., R.F., T.E., L.P. with help from F.L., S.P., G.M.C.

Performed experiments: J.K., R.F. with help from L.M.R., G.G., E.M.P., J.W.

Analyzed the data: J.K., R.F., T.V.

Supervised the project: L.P., G.M.C.

Wrote the manuscript: J.K., R.F., T.V., L.P with input from all authors

## Competing interest

G.M.C., J.K., L.P., R.F., T.E. filed a patent application on work presented here. G.C. is a co-founder of Editas Medicine and has other financial interests listed at: https://arep.med.harvard.edu/gmc/tech.html. T.V. has received PhD studentship funding from AstraZeneca.

## Methods

### Cell lines

HEK293T cells were acquired from AMS Biotechnology (AMS.EP-CL-0005) and the HAP1 *ΔMLH1* cell line was purchased from Horizon Discovery (HZGHC000343c022). HEK293T cells were cultured in DMEM (Invitrogen) and HAP1 cells in IMDM (Invitrogen), both supplemented with 10% FCS (Invitrogen), 2 mM glutamine (Invitrogen), 100 U ml^−1^ penicillin and 100 mg ml^−1^ streptomycin (Invitrogen) at 37 °C and 5% CO_2_.

### Compounds

Palbociclib (S1116 Selleck), Paclitaxel (Cambridge Bioscience), Doxycycline (Selleckchem), Pifithrin-alpha (Merck Life Science UK Limited, P4359-5MG) were dissolved in DMSO to generate stock solutions of 10 mM. Pyromycine (Thermo Fisher Scientific) was dissolved in water to a concentration of 10 mM and used at 2 µg/ml.

### pegRNA cloning

Regular pegRNAs for transient transfection were cloned using golden gate assembly as described in (*14*). Briefly, for each pegRNA, forward and reverse oligonucleotides were ordered for spacer, scaffold and 3’-extensions (IDT or Merck). The pegRNA acceptor plasmid (Addgene #132777) was linearized with BsaI, oligonucleotides hybridized, and the scaffold phosphorylated. The components were assembled using a golden gate reaction (with BsaI and T4 ligase) and transformed into XL10 gold ultracompetent bacteria (Agilent). Lentiviral engineered pegRNA plasmids were cloned the same way but using a lentiviral epegRNA acceptor construct as starting point (Supplementary Table 1) which was linearized with BsmBI instead of BsaI. In addition, an optimized scaffold was used (cr772, gtttaagagctaagctggaaacagcatagcaagtttaaataaggctagtccgttatcaactcgaaagagtggcaccgagtcggtg)

### Inserting loxP sites into LINE-1s by transient transfection

One day before transfection, 500,000 HEK293T cells were plated into each well of a six well plate and two cell pellets with 2 million cells each were frozen for RNA extraction. 2 μg of PE2 plasmid (Supplementary Table 1) and 500 ng of pegRNA plasmid (Supplementary Table 1) were transfected per well using Lipofectamine LTX (Invitrogen, using 2.5 µl plus reagent and 7.5 µl LTX solution). After 3 days, cells from 3 wells were frozen for RNA extraction. The remaining 3 wells were split 1:5 and replated in a 6 well plate. Remaining cells from the split were frozen for amplicon sequencing. The splits and cell pellet freezings were repeated at day 7, 9, and 11.

### Generating HAP1 *ΔMLH1* and HEK293T cell lines that stably express prime editor

HAP1 and HEK293T cell lines expressing prime editors were generated by cotransfecting pCMV-hyPBase (*60*) and pPB-TREG3G-PE2-rtTA3G-P2A-eGFP (Supplementary Table 1) (*61*). First, 500,000 HAP1 *ΔMLH1* and HEK293T cells were each seeded into one well of a six-well plate one day before transfection. For each transfection, 3 µg of each plasmid was mixed with 6 µl of Plus reagent and 7.5 µl of Lipofectamine LTX (Invitrogen) reagent, incubated for 30 min and then added to the cells. At two weeks post transfection, cells were sorted into single clones based on eGFP expression. To test prime editing on a single locus, we infected cells with a lentivirus containing a pegRNA targeting the HEK3 site and encoding a loxP sequence (Supplementary Table 1). We selected for guide integration with 2µg/ml puromycin and then induced prime editor expression with 1 µM doxycycline and determined editing rates by visualizing amplicons (Supplementary Table 2) on the bioanalyzer (Agilent).

### Lentivirus production

Lentivirus was produced in HEK293FT cells that were transfected with Lipofectamine LTX (Invitrogen). 0.9 μg of a lentiviral vector, 0.9 μg of psPax2 (Addgene 12260), 0.2 μg of pMD2.G (Addgene 12259) were mixed in 500 µl Opti-MEM together with 2 μl PLUS reagent and incubated for 5 min at room temperature. 6 μl of the LTX reagent was added and the mix which was incubated for another 30 min at room temperature. 500 µl of the transfection mix was then added to 80% confluent cells in 2 ml DMEM media in a well of a six well plate. After 48h the supernatant was collected and stored at -20°C. Fresh media was added to the cells and harvested 24h later. The two harvests were kept separate. For virus titration, Lenti-X GoStix Plus (Takara) was used following the manufacturer’s protocol.

### Flow cytometry

Samples were run on the CytoFLEX Flow Cytometer (Beckman) for analysis. The data was acquired with the CytExpert software and analyzed with FlowJo V10 or CytoExploreR. Events were first gated for cells based on forward and side scatter. Next, singlets were distinguished from doublets based on width and height of the side scatter light. Finally, cells were analyzed for their respective fluorescence channels. Sensitivity was set so that the mean fluorescence intensity of the negative population was around 10^1^ - 10^3^. Cells were sorted for single clones on either Sony MA900, Sony SH800S, or on MoFlo XDP for sorting.

### LINE-1 editing time course

500,000 HAP1 and HEK293T cells stably expressing piggyBac-integrated prime editor and pLenti-ePEG-LINE-1-loxPsym were seeded into 6 well plates and editing was induced with 1 µM doxycyclin in the presence of 10 µM PFT-alpha. The cells were split every 2 or 4 days and reseeded at approximately 30% confluency. The remaining cells were harvested for gRNA extraction and analysis.

### Maintenance and expansion of human iPSCs

Human induced pluripotent stem cells (iPSCs) were maintained in mTeSR medium on plates pre-treated with Matrigel from BD Biosciences. The iPSCs underwent regular subculturing, where they were exposed to TrypLE (Thermofisher # 12604013) for a duration of 5 minutes, followed by neutralization using an equivalent volume of mTeSR and subsequent centrifugation at 300 g for 5 minutes. The resulting iPSC clusters were mechanically dispersed to achieve a uniform cell suspension. This suspension was then seeded on plates coated with Matrigel, at a seeding density of 30,000 cells per square centimeter. The medium was enriched with 10 µM Y-27632 ROCK inhibitor (Ri) from Millipore (688001) during the initial 24 hours of culture. For single-cell cloning, 96-well plates were pre-treated with Matrigel (50µL per well). A specialized medium was created by mixing 10% CloneR™ (StemCell Technologies #05888) and 10 ng/l Pifithrin-α in mTeSR™ medium, which was then dispensed into the Matrigel-coated wells. Cells were sorted using fluorescence-activated cell sorting (FACS) into individual wells, each containing the pre-warmed cloning medium, at a seeding rate of one cell per well. To minimize disruption, the media in the plates was not replaced for the initial 48 hours.

### Maintenance and expansion of human skin fibroblasts

Human dermal fibroblasts (ATCC PCS-201-010) were maintained at 37°C with 5% CO_2_ and 5% O_2_ using low glucose DMEM (ThermoFisher 11885-084) enriched with 15% FBS (GenClone 25-550) and 1% Penicillin-Streptomycin (ThermoFisher 15140122).

### Editing of human skin fibroblasts

In the process of establishing prime editing lines, 200,000 cells underwent nucleofection with 100 ng of transposase (Super piggyBac Transposase - SystemBio PB210PA-1) and 500 ng of PiggyBac-PEmax. These procedures utilized the P2 Primary Cell 4D Nucleofector kit (Lonza V4SP-2096) and the 4D-Nucleofector system from Lonza, following the DS-150 program. Post-nucleofection, the cells were recovered at room temperature for 45 minutes before being seeded in a 24-well plate with 500 µL of DMEM media. Two days after nucleofection, cells were selected with 400 ng/mL Puromycin (ThermoFisher A1113802) for one week. Afterwards, surviving cells were transduced with the lentivirus epegRNA lentivirus, which included the addition of 6µg/mL of polybrene, followed by 5µg/mL of blasticidin selection two days post-transduction. Surviving cells were then single-cell cloned. Surviving clones were expended and editing efficiency was subsequently assessed as described below.

### Editing of human iPSCs (PGP1)

For this experimental procedure, a mixture of 2.5µg of PEmax and 1µg of Transposase was combined with 7.5µg of lipoSTEM. After two days, 1 µg/mL of puromycin was added to the culture. The cells were selected for two weeks. Afterwards, the cells were transduced with epegRNA lentivirus, which included the addition of 6µg/mL of polybrene, followed by 5µg/mL of blasticidin selection two days later. After 25 days of combined selection with puromycin and blasticidin, 400,000 cells were plated in a 6-well plate together with Ri, 1 µM doxycycline and 10% CloneR™ (StemCell Technologies #05888) in mTeSR medium, 10 ng/L Pifithrin-α, and maintained for 17 days. Surviving cells were then single-cell cloned. Surviving clones were expended and editing efficiency was subsequently assessed as described below.

### Long-term editing of a HEK293T cell line

HEK293T cells stably expressing epegRNA and PEmax were grown with DMEM 10% FBS and 2µg/mL of puromycin, 5µg/mL, and 1µM of doxycycline over the course of a year. Cells were single-cell cloned every 3-4 months and surviving clones were analyzed with NGS. The clone with the highest editing rate was expanded. This process was repeated over the course of 1 year. After one year of editing, the clone with the highest editing rate was analyzed with nanopore sequencing (ligation and Cas9 sequencing).

### Amplicon sequencing of LINE-1s

Genomic DNA was either extracted from cell pellets using the DNeasy Blood & Tissue kit (Qiagen) with addition of 50 µg RNaseA in the pellet resuspension step or cell pellets were prepared by direct lysis using home-made quick extract buffer (1 mM CaCl_2_, 3 mM MgCl_2_, 1 mM EDTA, 1% Triton X-100, 10 mM Tris pH 7.5) with freshly added proteinase K (0.2 mg/ml) followed by 15 min incubation at 65°C and 20 min incubation at 95°C. For library preparation, 3 µl of genomic DNA (∼ 30 ng) or 3 µl of direct lysis extract were used as template in 50 µl PCR reactions using KAPA HiFi HotStart ReadyMix (Roche), primers P3/4 and P5 (Supplementary Table 2). This first PCR was ran for 16-19 cycles (3min 95°C, 18x(20sec 98°C, 15sec 66°C, 30sec 72°C), 5min 72°C) and then purified with Agencourt AMPure XP beads in 1:1 ratio (beads to PCR reaction volume). Sequencing adaptors and barcodes were added with a second round of PCR using the KAPA HiFi HotStart ReadyMix (Roche), primers P6 and P7 (Supplementary Table 2), and 1 µl of the purified first PCR product as template. Amplicons were purified with Agencourt AMPure XP beads in 1:1 ratio (beads to PCR reaction volume) and quantified using the Nanodrop 2000. The amplicons were pooled together and sequenced on the Illumina MiSeq 2500 (500 cycles, 250 paired-end).

### Scramble time course with TAT-Cre protein

Without Cre reporter: HAP1 cells with 301 monoclonal loxPsym site integrations (HAP1-loxPsym(301)) were plated in 6 well plates with 1.5 million cells per well in 1.5 ml of IMDM medium supplemented with FBS. 5 µM or 2 µM of TAT-Cre protein (Cambridge Biochemisty Department) was added to the cells. TAT-Cre containing medium was removed 4 hours after induction and replaced with 2 ml of fresh medium. Cells were trypsinized and seeded into T150 flasks after 16 hours and harvested and split at the indicated time points.

With Cre reporter: HAP1 cells with 301 monoclonal and HEK293T cells with 638 monoallelic loxPsym site integrations and expressing the tBFP-Cre reporter construct (HAP1-loxPsym(301)R and HEK293T-loxPsym-638R) were plated in 6 well plates with 1.5 million cells per well in 1.5 ml of IMDM medium without FBS. 1 µM TAT-Cre protein was added to the cells 30 minutes after seeding and removed 4 hours later. 2 ml of fresh medium containing FBS was added to the cells. The cells were trypsinized and seeded into T150 flasks after 16 hours. The cells were harvested and split at the indicated time points and the fraction of BFP-positive cells was monitored by flow cytometry. HEK293T-loxPsym-638R and HAP1-loxPsym(301)R cells were sorted for BFP-positive cells at day 13 and day 15 respectively.

### Cell cycle inhibition

500,000 HAP1-loxPsym(301) cells were plated in each well of a 6 well plate. One day later, 3 wells were treated with 500 nM palbociclib for 24 hours. To confirm cell cycle inhibition, one well of cells was harvested, stained for DNA (Hoechst-33342) and analysed using flow cytometry. For the remaining wells, the medium was replaced with IMDM without FBS. After 1 hour, 1 µM TAT-Cre protein was added to the cells and removed 4 hours later. Palbociclib was kept in the media throughout the process. To remove extracellular Cre, cells were incubated in TrypLE Express (Theromfisher) for 10 minutes. Fresh media containing Palbociclib was added to the cells. 16 hours later, the cells were trpysinized. 2 million cells were harvested for long-read sequencing and the rest was plated in larger flasks and grown in full media without Palbociclib to restart the cell cycle.

### High molecular weight DNA extraction

Between 2 and 4 million cells were harvested by centrifugation and high molecular weight (HMW) genomic DNA was extracted using the Monarch High Molecular Weight DNA Extraction kit (NEB) and agitation speeds of 1,500 rpm.

### Long read whole-genome sequencing

HMW DNA was sheared to 20 kb using Covaris G-Tubes (for shallow-depth sequencing with MinION) or using the Megaruptor 3 (Diagenode; for high-depth sequencing with PromethION). Nanopore sequencing libraries were prepared using the Ligation Sequencing Kit V14 (Oxford Nanopore) and sequenced on the MinION Mk1B using R10.4.1 flow cells (FLO-MIN114) or on the PromethION. Base calling was performed using the ‘Super high accuracy model’ (dna_r10.4.1_e8.2_400bps_sup.cfg) of the guppy basecaller (versions 6.3.8 or 6.4.6).

### Identification of insertion sites for loxPsym sites

Long-read whole genome sequencing fastq files were aligned to the hg38 reference genome using minimap2 (*62*, *63*). The resulting sam files were sorted and compressed into bam files using samtools (*64*). Structural variants were called with relaxed settings using Sniffles2 (*65*) with the --non-germline --phase --output-rnames --tandem-repeats --minsupport 1 --minsvlen 25 parameters. Custom R scripts (Github: https://github.com/jonas-koeppel/genome_scramble) were used to identify loxPsym insertion sites and call Cre-induced rearrangements. For the identification of insertion sites, R::agrep with the max.distance = 0.2 option was used to find variants that contain ‘TAACTTCGTATAATGTACATTATACGAAGTTA’ in the ALT sequence and subsequently filtered for insertions < 50 nt. Clonal insertions in HAP1 were defined as sites on standard chromosomes with >5 supporting reads and an allele frequency of > 0.5. Clonal insertions in HEK293T were defined as sites on standard chromosomes with at least 3 supporting reads and an allele frequency of > 0.1. Insertion sites are deposited as Data S1.

### Identification of Cre-induced variants

Raw structural variants were called using nanomonsv (*66*). The unscrambled parentals were considered as control and the scrambled cell lines as tumor. To call variants in the unscrambled negative controls, unedited parental HAP1/HEK293T cells were designated control. One supporting read was required for the tumor and 0 for the control and a panel of normals. The raw structural variants were filtered to (i) contain fragments of a loxPsym site at the break junction (“TAACTTCGTAT | ATACGAAGTTA | AATGTACATTAT | GTATAATGTAC”) (ii) Start and end within 300 bp of a nicking site of the LINE-1 targeting pegRNA (allowing 3 mismatches to the protospacer). (iii) Do not have coverages higher than 5x the average sequencing depth (regions with higher depth are usually ambiguously mapped). (iv) Are located on chromosomes 1-22 + X, Y. A schematic of the variant calling process is shown in Supplementary Figure 7. The filtered variant calls are deposited as Data S2.

### Identification of variants in edited clones

To identify variants associated with the loxPsym insertion process (Supplementary Figure 14), but unrelated to scrambling. Raw variant call data for unscrambled cells were filtered for > 5 reads and an allele frequency > 0.5 (HAP1) or > 3 reads and an allele frequency > 0.1 (HEK293T). Insertions were categorized as loxPsym if the insertion sequence contained any of the following sequence: TAACTTCGTAT, ATACGAAGTTA, or AATGTACATTAT, as pegRNA if they included ‘GTACATTATACGAAGTTATAGGAGGAACTGGTACCATTCCTTCTGAAACTAT’ or as prime editor for ‘TTAACCCTAGAAAGAT’.

### Further genome analysis

Coverages were estimated using samtools bedcov, considering reads with at least quality 10 and binning in 10 kb windows. For plotting, depth was calculated using mosdepth (0.3.3 (*67*)) and 50 kb binning windows (or 500 kb windows for Supplementary Figure 14). B allele frequencies were calculated using amber in tumor only mode (*68*). The 4DNFI1E6NJQJ HiC data set was used for HAP1 cells and visualized using plotgardner (*69*).

### Simulations of contact frequency and 3D distance

A matrix of all possible recombinations was generated by simulating rearrangements for all pairs of loxPsym sites on each chromosome. In the naive model each variant was equally likely. The observed rearrangements were then compared to all possible ones. For the exponential decay model, the simulated rearrangements were adjusted to decrease in frequency of occurrence with increased distance (between 20% and 0.1% per Mb in 0.1% increments). The distributions resulting from simulations were compared to the observed data and their Wasserstein distance calculated. Pearson correlations between sequencing reads (0 if no rearrangement between a given pair was observed) and the logarithms of variant length or Hi-C read pair numbers (4D nucleome, 4DNFI1E6NJQJ) were estimated using a linear model.

### Cas9-enrichment long-read sequencing of edited LINE-1s

Cas9-enrichment was performed according to the Cas9 sequencing kit protocol (CAS9106 Protocolv109, Oxford Nanopore Technologies). Briefly, HMW DNA was sheared to 20 kb using Covaris G-Tubes and 5 µg were dephosphorylated. ALT-R tracrRNA and crRNA targeting LINE-1 with a loxPsym site insertion (ACATTATACGAAGTTATAGG) were ordered from Integrated DNA technologies (IDT) and complexed with HifiCas9 V3 (IDT) to form RNPs. Dephosphorylated DNA was treated with Cas9 RNPs for 1 hour at 37dC. Sequencing adapters were ligated to the cut DNA. The ligation step was extended to 1 hour at room temperature. The libraries were then purified using Ampure XP beads (Agilent), washed with Long fragment buffer and eluted in elution buffer for 1 hour. The libraries were sequenced with the MinION Mk1B using R9.4.1 flow cells (FLO-MIN106). Base calling was performed using the ‘Super high accuracy model’ (dna_r10.4.1_e8.2_400bps_sup.cfg) of the guppy basecaller (versions 6.3.8 or 6.4.6). The resulting read files were filtered with seqkit to only contain reads that cover LINE-1s by matching the following sequence (“AGGAGGAACTGGTACCATTCCTTCTGAAACTATT”) while allowing up to 3 mismatches. The filtered read file was aligned to the hg38 reference genome (Dec. 2013 GRCh38/hg38) using minimap2. The number of reads covering each LINE-1s with up to two mismatches from the protospacer (Data S3) were then determined using bedtools multicov (*70*) requiring a mapping quality score of at least 5.

### Chromatin states and epigenetic analyses

The following publicly available datasets were collected: HEK293T: DNase-seq ENCFF969MBJ, H3K4me1-ChIP-seq SRR10981645, H3K36me3-ChIP-seq SRR5627148, H3K9me3-ChIP-seq SRR11453034, H3K27me3-ChIP-seq SRR8937480, H3K4me3-ChIP-seq SRR8937479, and H3K27ac-ChIP-seq SRR1016003. HAP1: DNase-seq ENCFF162WTC, H3K4me1-ChIP-seq ENCFF639UYT, H3K36me3-ChIP-seq ENCFF216JJJ, H3K9me3-ChIP-seq ENCFF528UHF, H3K27me3-ChIP-seq ENCFF708HAB, H3K4me3-ChIP-seq ENCFF461TZF, and H3K27ac-ChIP-seq ENCFF742SZS (Supplementary Table 3) (*71*, *72*). For data sets from ENCODE, hg38 aligned bam files were downloaded and for data sets from the sequence read archive, fastq files were downloaded and aligned to the hg38 genome build using bwa-mem. The resulting bam files were binarized using ChromHMM and a 15-state model was learned (Supplementary Figure 6, Data S4). In addition the GSM4625025 for CTCF-ChIP-seq in HAP1 and ENCSR000DTW for CTCF-ChIP-seq in HEK293T data sets were visualized in IGV.

For the comparison of epigenetic states in edited (> 1 read in the Cas9 sequencing experiment) and non-edited LINE-1s (0 reads in the Cas9 sequencing experiment), the fraction of nucleotides in each ChromHMM state was calculated across a sequence window consisting of the LINE-1 and 3 kb of flanking sequence (to mitigate possible misalignment of short read data in repetitive LINE-1s). States that were found in less than 10 edited LINE-1s were filtered out. Enrichments (risk ratios) were calculated by dividing the fraction in each state across edited and non-edited LINE-1s.

To compare the epigenetic states of clustered edited and non-edited LINE-1s, we intersected LINE-1s including 2 kb, 5 kb, 10 kb and 15 kb upstream and downstream with the chromHMM states from HAP1 and HEK293T cells. For each element, we calculated the fractional overlap across all states and conducted k-means clustering with 8-10 clusters to group elements that had similar epigenetic modifications. Clusters where elements had a mean fractional overlap > 0.75 for a state were annotated with this state, clusters where states had a mean fractional overlap ≤ 0.75 and > 0.25 were annotated using the top two states, and clusters where all states had a fractional overlap ≤ 0.25 were labeled as unassigned. Each LINE-1 was assigned to a bin based on the number of supporting reads.

### RNA sequencing

RNA was extracted from 2-5 million flash frozen cells using the RNeasy plus mini kit (Qiagen). Libraries were prepared using the NEBNext® Ultra™ II Directional RNA Library Prep Kit for Illumina (New Englands Biolabs), multiplexed, and sequenced on two Illumina-HTP Novaseq 6000 lanes using 150 bp paired end reads. The median insert size was 280 bp (quartiles 231, 355). Between 58 and 191 million reads were generated per sample. Salmon (2.0.0 (*73*)) was used to quantify transcripts against a salmon index built for the hg38 human reference genome. Further analysis was done using custom R scripts. Transcripts were collapsed to gene level using tximport (1.22.0) (Data S5). Normalization (rlog) and differential expression analysis was performed using DESeq2 (1.34.0). The ensembl v110 release was used for gene structure annotation (imported into R using biomaRt (2.50.3). For all analyses involving genes, the gene lists were filtered to contain only “protein_coding”, “lncRNA”, “snRNA”, “snoRNA”, “miRNA”, “rRNA”, “ribozyme” genes with expression base means > 20. Gene set enrichment analysis was done using fgsea (1.20.0) and the h.all.v2022.1.Hs.symbols.gmt gene set (https://data.broadinstitute.org/gsea-msigdb/msigdb/release/2022.1.Hs/)

### Features of structural variants

To understand the selection effects acting on structural variants we conducted an enrichment analysis across 49 features comparing variants that were observed at early and late time points. All features are described in Supplementary Table 4. PhyloP conservation scores were downloaded from UCSC genome browser (*74*) and each structural variant was annotated with the mean, median and max across all overlapping bases. The fraction of conserved elements overlapping each structural variant was computed using GERP++ conserved elements (*75*). The fraction of each structural variant overlapping human accelerated regions was computed using annotations from Girskis et al. (*76*). Constraint z-scores passing all quality control checks for coding and non-coding regions were downloaded from gnomAD (*77*). Structural variants were annotated with the maximum, median and mean gnomAD z-score across all overlapping 1 kb windows as well as the fraction of windows with a z-score greater than 2, 3, and 4. Gene features (exon, 5’ UTR, 3’ UTR, stop codon, start codon, CDS and gene) were computed as the fraction overlapping each structural variant using GENCODE release v44 (*78*). The fraction of each structural variant overlapping super enhancers was calculated using annotations from SEdb 2.0 (*79*). The fractional overlap of each structural variant over TAD boundaries, A/B compartments and lamina-associated domains (LMNB1) from HAP1 cells was calculated using data from the 4D nucleome project (*80–82*). The fraction of the structural variant overlapping CpG islands was computed using annotations from the UCSC genome browser (*83*).

Candidate cis-regulatory elements in HAP1 cells were downloaded from ENCODE and for each variant the fractional overlap over each type of element was computed (*2*). The fractional overlap over chromatin states was computed using the HAP1 chromHMM annotation described previously. Finally, length of the structural variant was included as a feature. We tested for enrichment for deletions and inversions separately, normalizing each feature within each variant class. We performed logistic regression modeling whether the structural variant was observed at an early or late time point as a function of each feature individually. 95% confidence intervals for the fitted parameters were calculated using the standard normal distribution.

### HAP1 essentiality screen

The library and general screening procedure were described in (*30*). The screen in HAP1 cells is briefly described here. 20M Hap1-Cas+ cells were transduced in two biological replicates using a spinfection in 6 well plates with lentivirus aiming for a MOI of 0.3. The spinfection was carried out at room temperature for 30 minutes spinning at 1000g and the cells were supplemented with 8 μg/ml polybrene (hexadimethrine bromide, Sigma). 2 µg/ml puromycin was added to the cells 24 hours post infection. 2 µg/ml puromycin was kept in the media throughout the screen. Cells were grown for 14 days and time points were taken at days 3, 7, 10 and 14 post infection. For each time point at least 24 million cells were harvested. To derive relative fitness scores, the fold changes of guide abundance in sequencing reads between day 14 and the plasmid library were calculated for six independent guides per gene and collapsed to gene level by taking the average. Finally, the two biological replicates were merged by taking the average gene-level fold change (Data S6).

### Isolation of scrambled clones

13 days after scrambling (see section Scramble time course with TAT-Cre protein), Cre reporter expressing cells were single-cell sorted for tBFP expression (indicating successful Cre recombination) and haploidy (based on cell size in forward and side scatter) into 96-well plates containing 200 µl of growth medium per well. Cells were transferred to flasks after visible colonies were formed. High molecular weight DNA was extracted as above. Clones were sequenced at 3-5x whole genome coverage either on the minion or after on the PromethION after pooling the DNA from various clones.

### Competition assay between scrambled clones and parentals

Scrambled clones (tBFP-positive from the rearranged Cre reporter) were mixed with parental cells that do not express the Cre-reporter (tBFP-negative) in a 50:50 ratio and grown for 7 days. The cell ratio of parental to scrambled cells was determined on the indicated days by flow cytometry for tBFP expression.

### G-band karyotyping

Complex G-band karyotyping was performed and analyzed by Karyologic, Inc from samples of cryopreserved cells.

### Data availability

Processed data are deposited alongside this manuscript as Data S1-Data S6. Scripts used to analyze the data and additional necessary input files to run the scripts are deposited on https://github.com/jonas-koeppel/genome_scramble.

### Software

Genomics: bwa (0.7.17), bedtools (v2.29.0), ChromHMM (1.24), FlowJo (v10), guppy (6.3.8 and 6.4.6), minimap2 (2.22), mosdepth (0.3.3), sniffles2 (2.0.7), samtools (1.14), salmon, seqkit (v.2.0.0), IGV (2.16.2), R (4.1.3), nanomonsv (0.6.0) R packages: biomaRt (2.50.3), circlize (0.4.15), ComplexHeatmap (2.10.0), CytoExploreR (1.1.0), CytExpert, DESeq2 (1.34.0), fgsea (1.20.0), ggrepel (0.9.3), ggplot2 (3.4.4), ggpointdensity (0.1.0), plotgardnerer (1.4.2), plyranges (1.14.0), RColorBrewer (1.1.3), Repitools (1.40.0), ShortRead (1.46.0), spgs (1.0-3), StructalVariantAnnotation (1.10.1), tidyverse (1.3.2), VariantAnnotation (1.40.0), viridis (0.6.2), tximport (1.22.0) Python packages: numpy (1.21.6), pandas (1.2.3), pybedtools (0.9.0), pybigwig (0.3.13), pysam (0.16.0), python (3.7.7), sklearn (0.23.2), statsmodels (0.13.2)

## Supplementary Figures

**Supplementary Figure 1.**
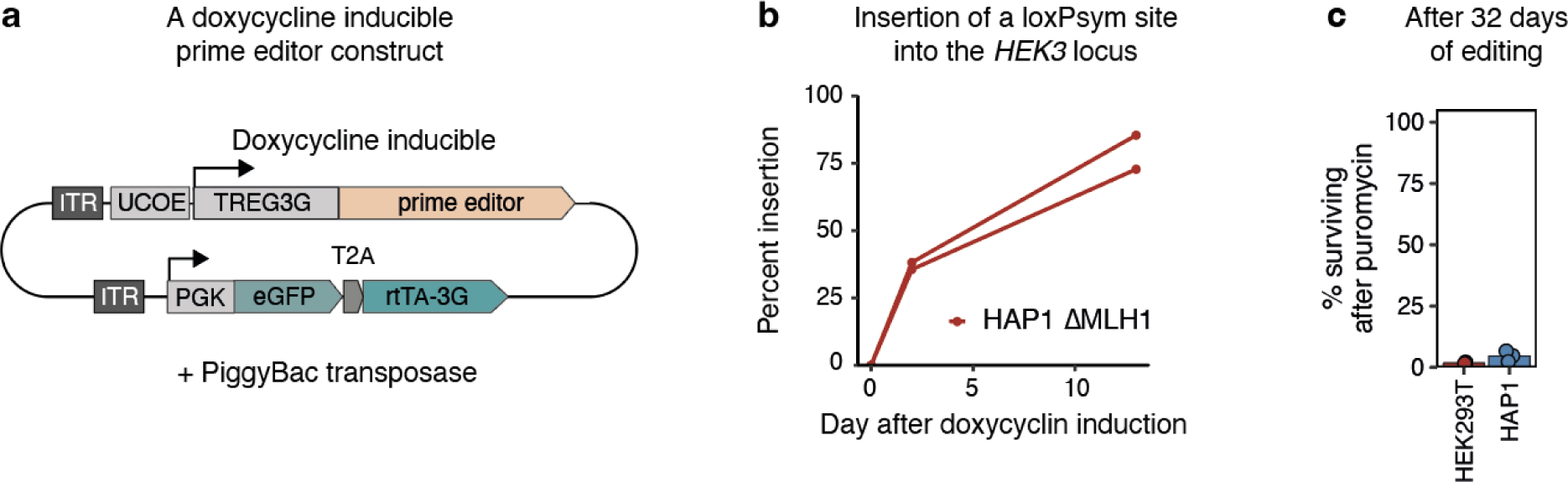
Highly efficient integration of loxPsym sites. (**A**) Schematic of the doxycycline-inducible prime editor expression construct on a PiggyBac transposon. UCOE: Universal chromatin opening element. TREG3G: Doxycycline-inducible promoter. rtTA-3G: transactivator protein. ITR: Inverted terminal repeats. (**B**) Integration efficiencies (y-axis) for a loxP site into the HEK3 locus over the course of 13 days (x-axis) in two clones (markers and lines). (**C**) The percent of surviving cells after puromycin treatment (y-axis) for HEK293T and HAP1 cells (x-axis) after 25 days of loxPsym site insertion into LINE-1s. The integrated pegRNA vector encodes for puromycin resistance.

**Supplementary Figure 2.**
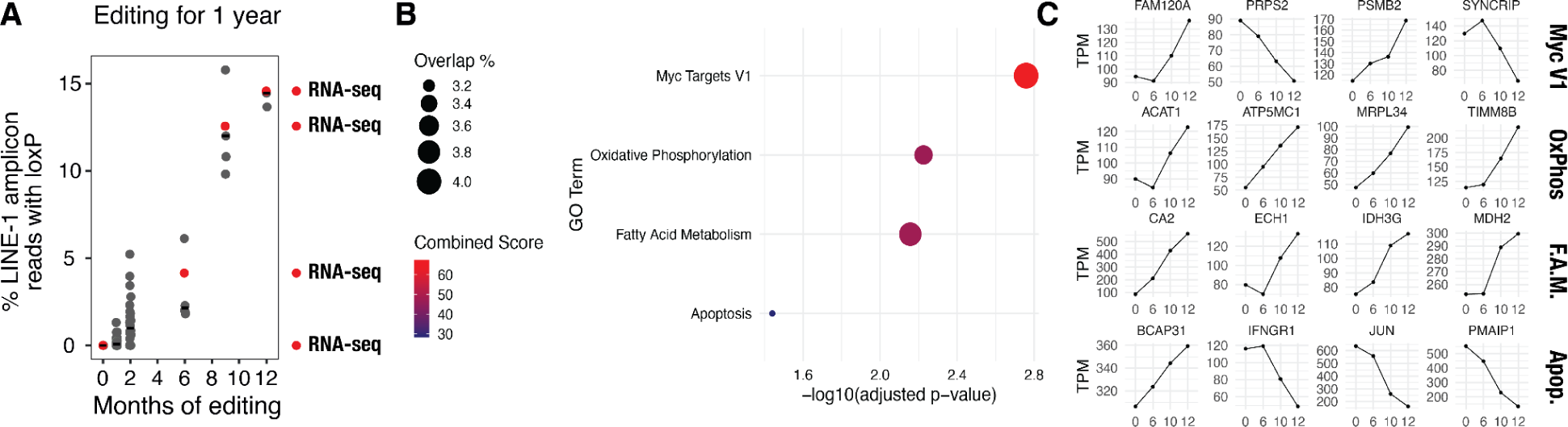
RNA-seq analysis of long-term edited HEK293 cells. (**A**) RNA-seq was performed at four different time points: 0, 5, 10, and 12 months (red dots). (**B**) GO term enrichment analysis of genes differentially regulated throughout the long-term editing process. Linear regression identified top-regulated genes using an absolute slope greater than 2 and R² value superior to 0.9. **(C)** Individual plots of four significantly regulated genes for each identified pathway. (Myc V1: Myc Targets V1; OxPhos: Oxidative Phosphorylation; F.A.M.: Fatty Acid Metabolism; Apop.: Apoptosis).

**Supplementary Figure 3.**
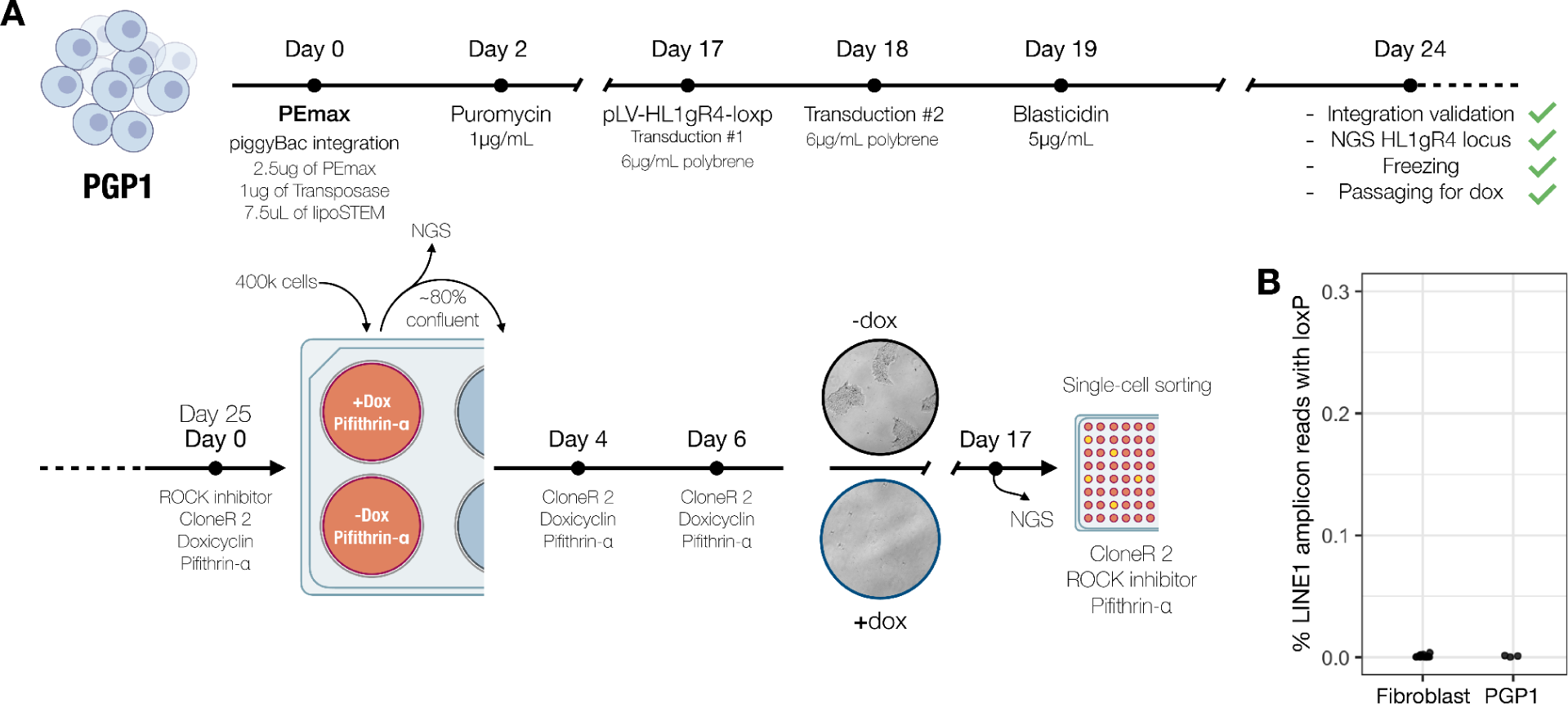
Editing of repetitive elements in human stem cells and skin fibroblast. (**A**) Graphical outline of the editing time course for human iPSCs PGP1. (**B**) Frequencies of LINE-1 amplicon reads with loxPsym insertions for Fibroblast and iPSCs (PGP1) after multiple rounds of subcloning.

**Supplementary Figure 4.**
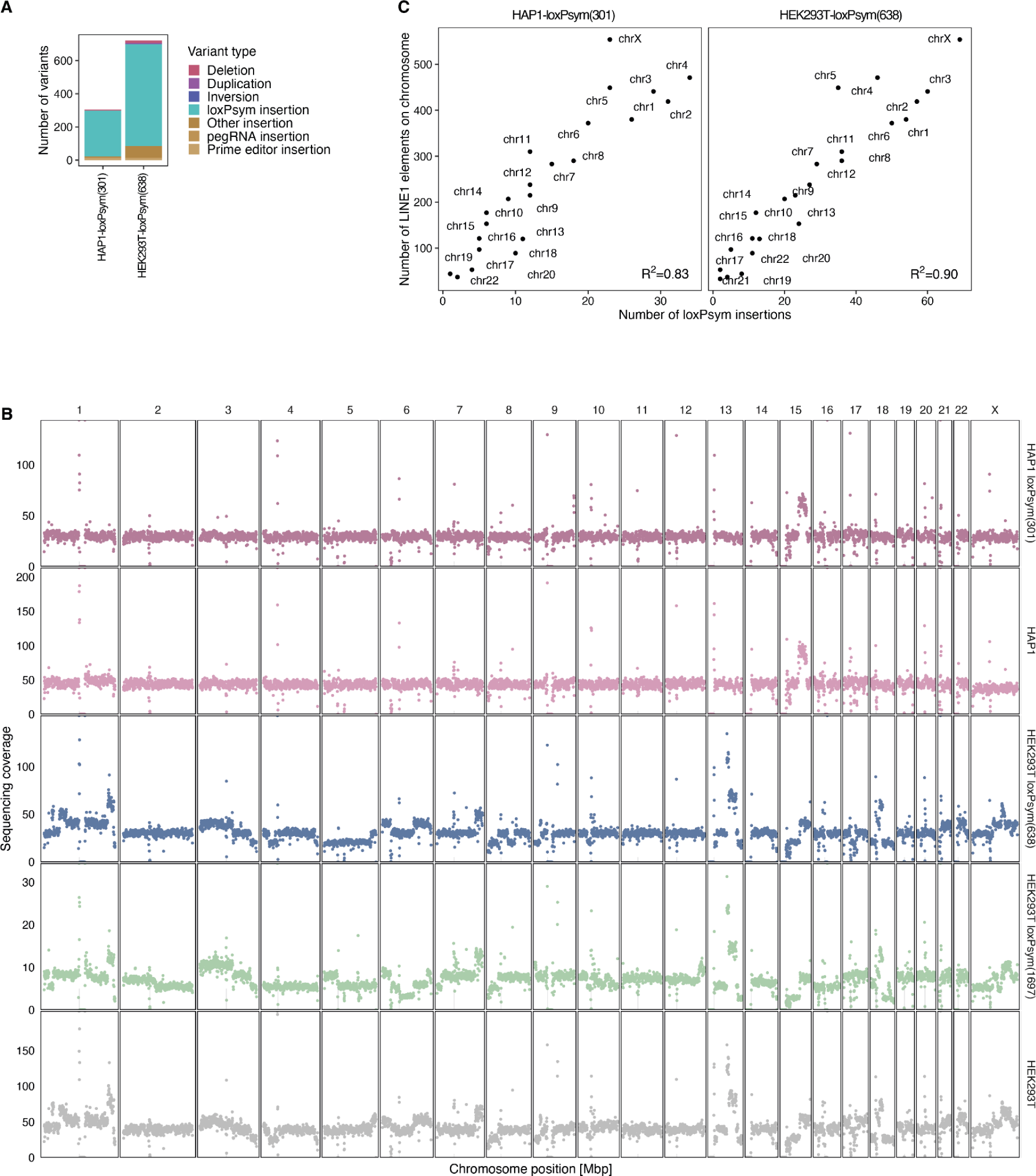
The number of loxPsym insertions is correlated with the number of LINE-1s per chromosome. (**A**) The number of structural variants (y-axis) for whole-genome sequenced clones (x-axis) colored by variant type. Only variants that are not also present in parental cells are shown. (**B**) Mean sequencing coverage (y-axis) across 500 kb bins (markers, x-axis) separated by chromosomes and clones (panels). (**C**) The number of loxPsym insertions (x-axis) versus the number of LINE-1 elements with up to two mismatches to the protospacer (y-axis) in HAP1 and HEK293T cells (panels) for chromosomes (markers).

**Supplementary Figure 5.**
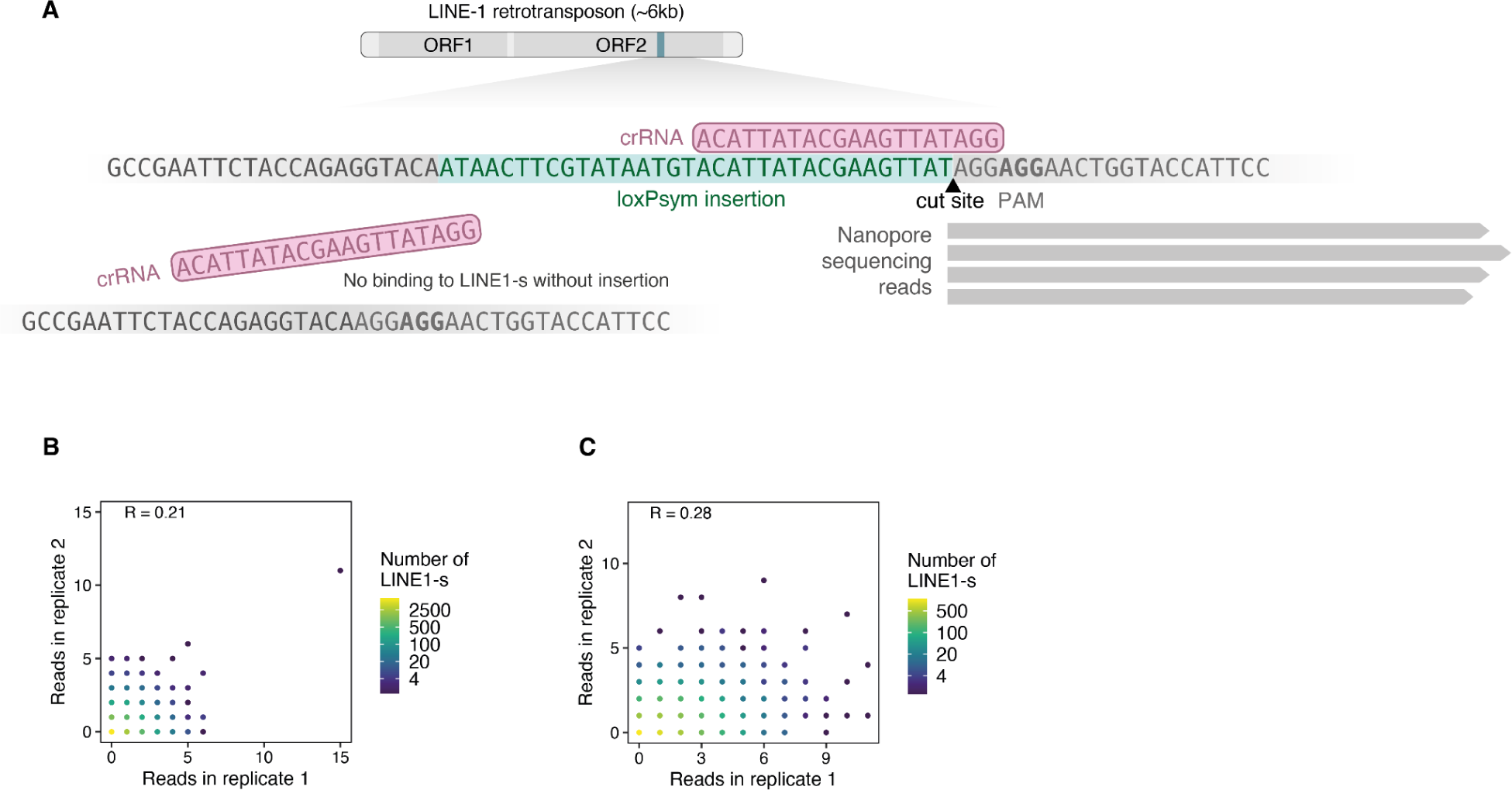
Sequencing with Cas9 enrichment to identify LINE-1s with loxPsym insertions. (**A**) Schematic of a strategy to map edited LINE-1s at high throughput. The sequence at the top represents a LINE-1 with loxPsym insertion (teal) to which the Cas9-enrichment crRNA can bind (pink). (**B**) The number of sequencing reads for each LINE-1 (marker) with up to 2 mismatches to the protospacer for two biological replicates (x- and y-axis) after 25 days of loxPsym site insertion into LINE-1s in HAP1 cells. Markers are colored based on the number of elements with that read number profile. (**C**) As in (B) but for HEK293T cells.

**Supplementary Figure 6.**
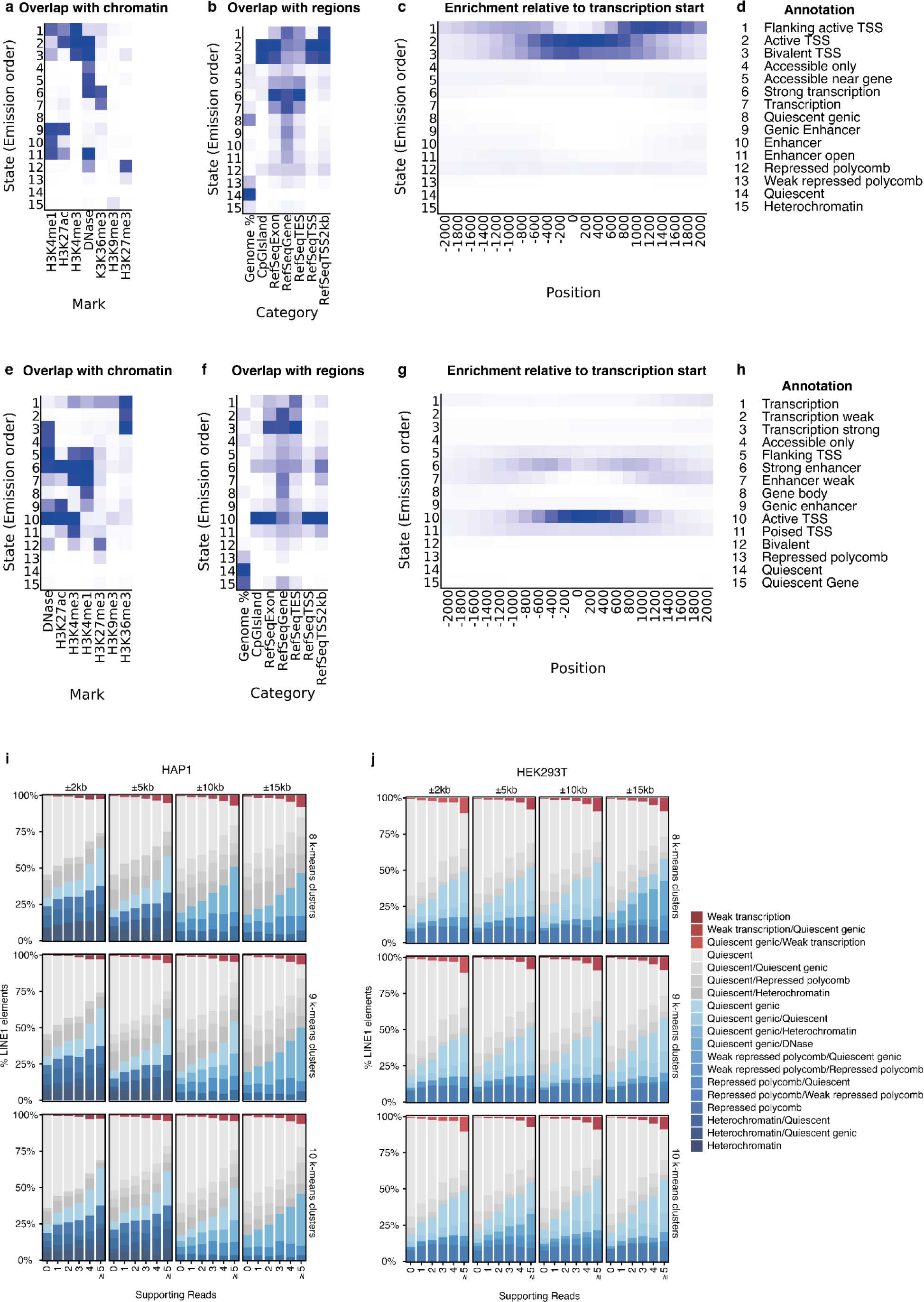
Chromatin states in HAP1 and HEK293T cells. (**A**) The overlap of ChromHMM emission states (y-axis) with chromatin marks (x-axis). (**B**) As in (A) but for different types of sequence regions in the genome. (**C**) The enrichment of ChromHMM emission states (y-axis) relative to varying distances to transcription start sites (x-axis). (**D**) Manual annotation of emission states based on (A-C). (**E-H**) As in (A-D) but for HEK293T cells. (**I**) The percent of LINE-1s (y-axis) in various epigenetic states (colors) stratified by the number of nanopore sequencing reads (x-axis) for k-means clustering with 8-10 clusters and considering 2, 5, or 10kb of sequence up-and downstream of the LINE-1. HAP1 cells. (**J**) As (I) but for HEK293T cells.

**Supplementary Figure 7.**
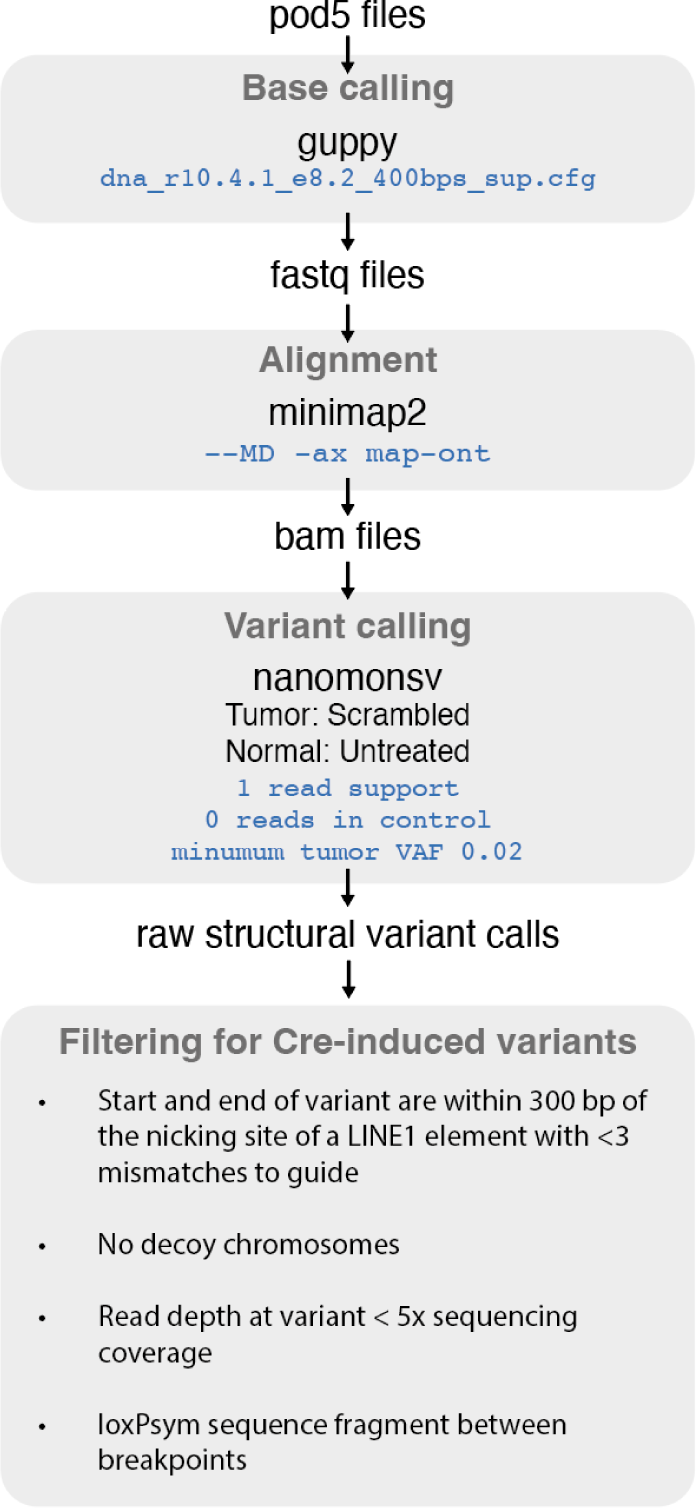
Pipeline for calling Cre-induced variants. Schematic outlining the bioinformatics pipeline used to process sequencing data and call Cre-induced structural variants. pod5 files were base called using Guppy and aligned with Minimap2. Nanomonsv was used with relaxed parameters to call raw structural variants which were filtered based on proximity to LINE-1s, absence of decoy chromosomes, read depth thresholds, and the presence of loxPsym sequence fragments between breakpoints.

**Supplementary Figure 8.**
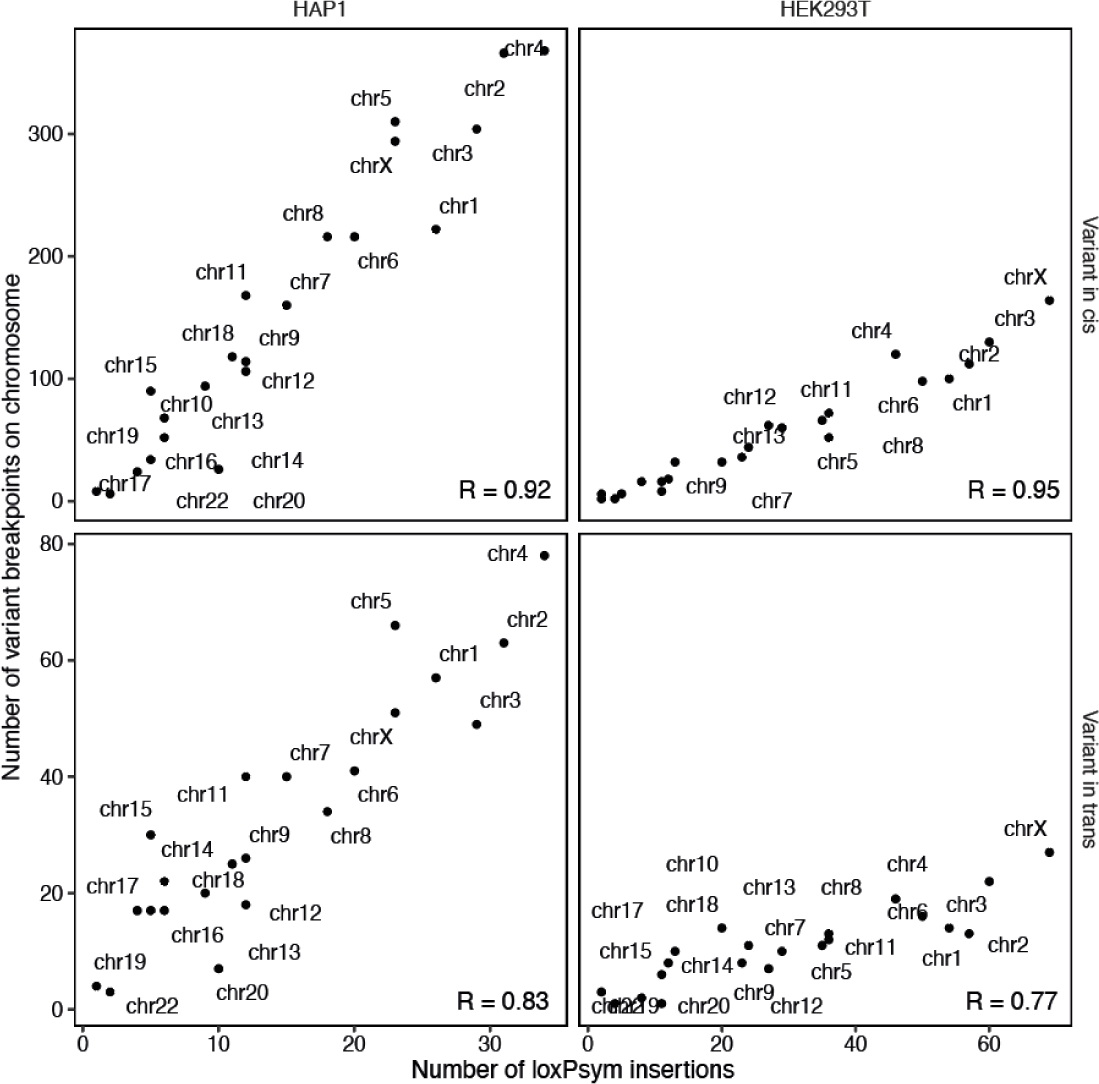
Patterns of Cre rearrangement. The number of loxPsym insertions (x-axis) versus the number of variant breakpoints on chromosomes (y-axis) in HAP1 and HEK293T cells (panels) for chromosomes (markers).

**Supplementary Figure 9.**
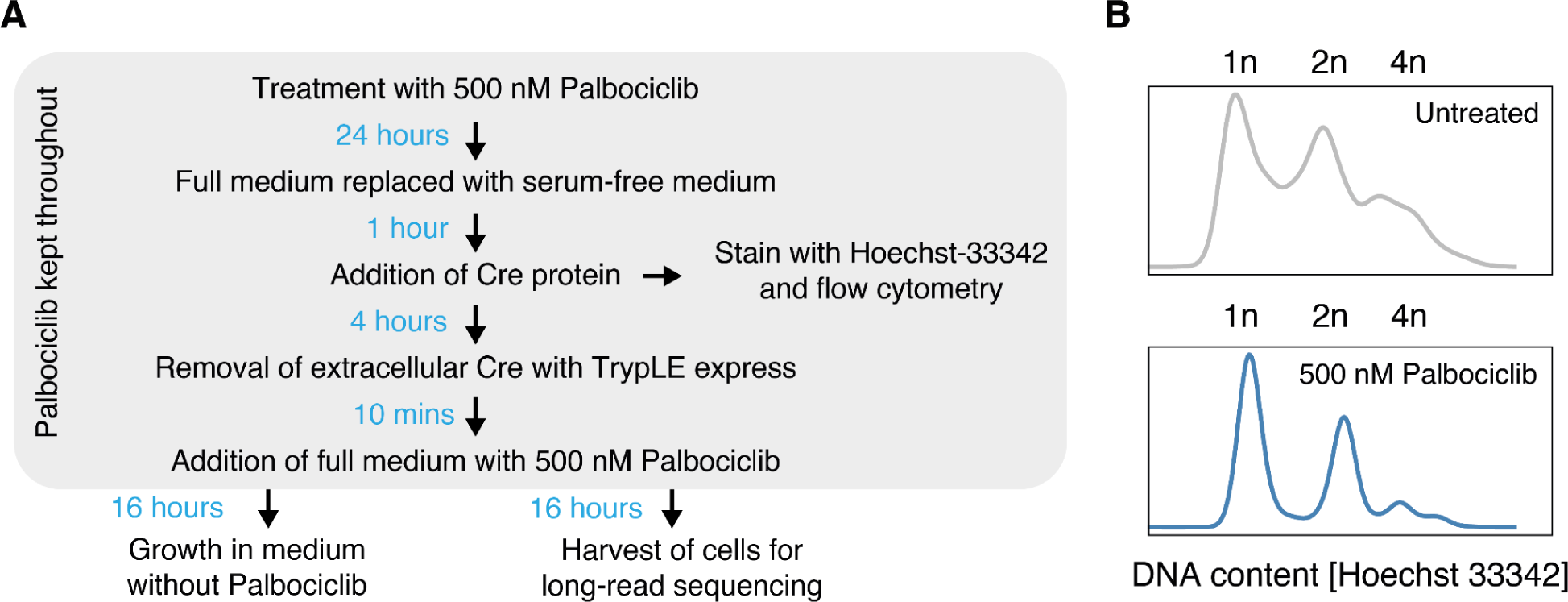
Scramble with cell cycle inhibition. (**A**) Schematic illustrating the experimental protocol for treating cells with palbociclib. (**B**) Frequencies (y-axis) of DNA content (x-axis) in untreated cells (top panel) and cells treated with 500 nM palbociclib (bottom panel). Likely genome copies (1n, 2n, 4n) are indicated.

**Supplementary Figure 10.**
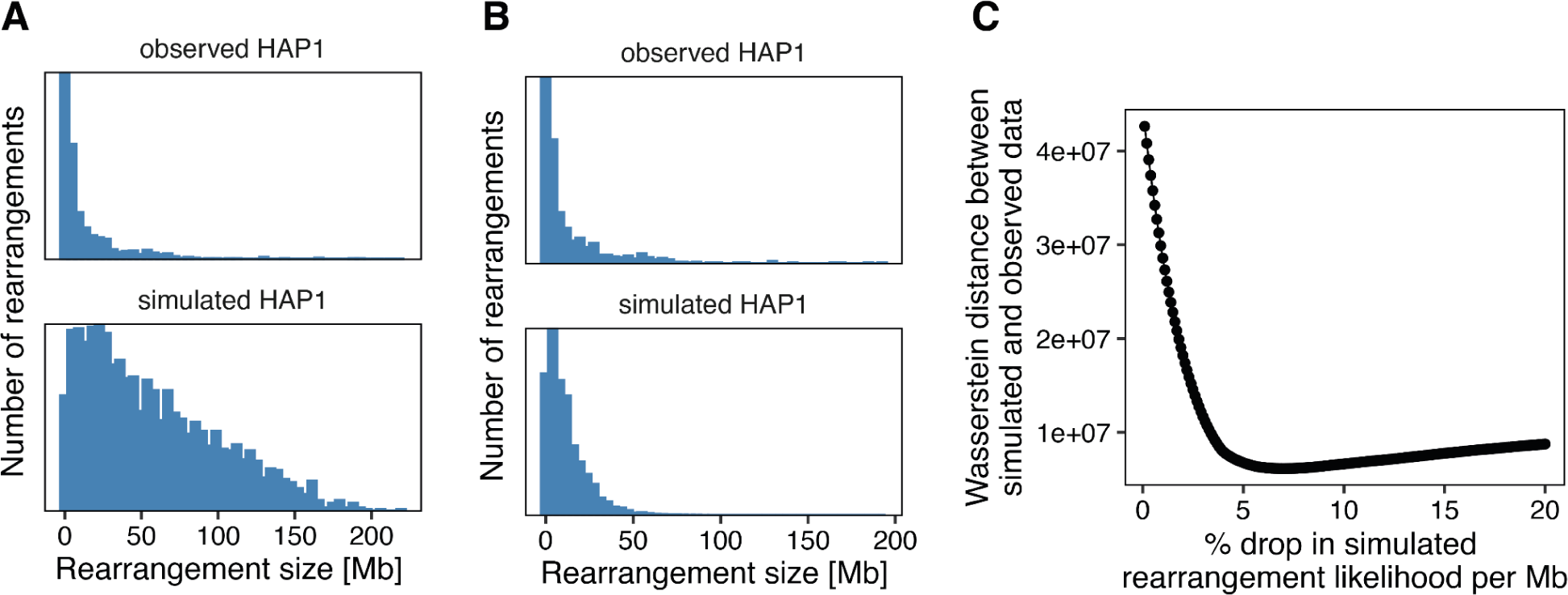
Simulated and observed deletions and inversions. (**A**) Histograms contrasting the number of observed rearrangements (upper panel, y-axis) with those generated by simulation assuming equal likelihood of recombination between any two sites on the same chromosome (lower panel) across different rearrangement sizes (x-axis, with 50 bins) in HAP1 cells. (**B**) As (A) but with a model assuming a 7.5% decrease in recombination likelihood per megabase (Mb). (**C**) The Wasserstein distance between simulated and observed data (y-axis) for various reductions in rearrangement likelihood per Mb (in increments of 0.1%, x-axis).

**Supplementary Figure 11.**
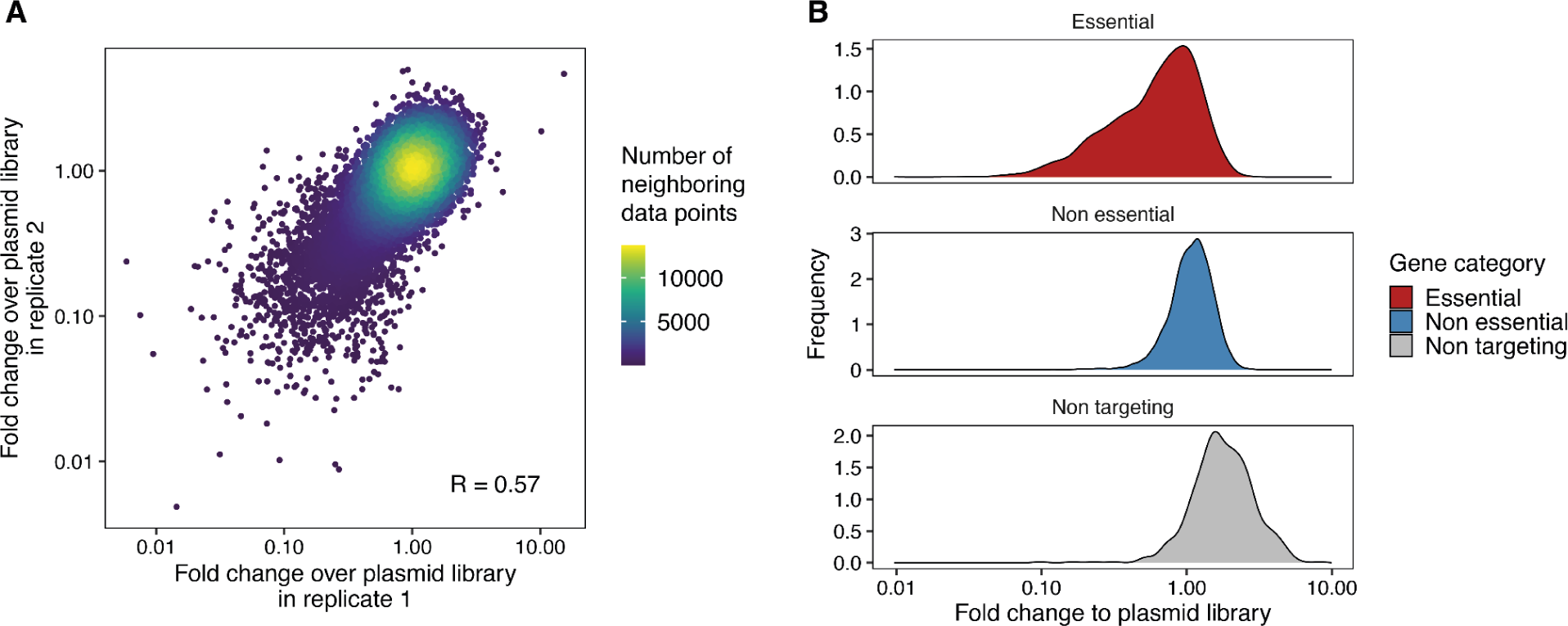
A genome-wide CRISPR screen in HAP1 cells. (**A**) Fold change in gene perturbation frequency over the plasmid library between two independent replicates (axes) colored by the number of neighboring data points. (**B**) Frequencies (y-axis) of fold change to plasmid library (x-axis) for gene perturbations categorized by gene essentiality (panels and colors).

**Supplementary Figure 12.**
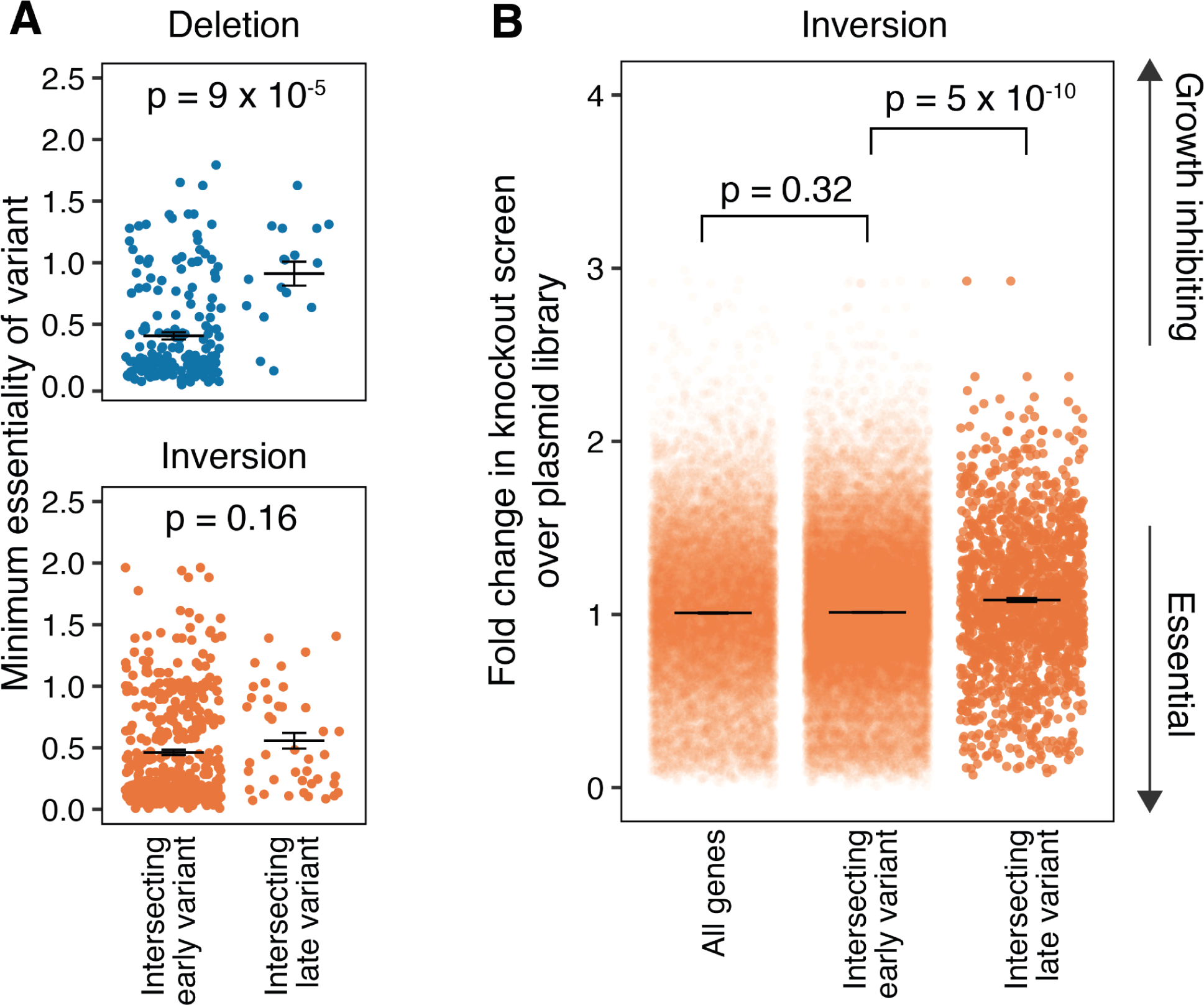
Essentiality of genes intersecting deletions and inversions. (**A**) Median essentiality scores for genes (y-axis) affected by deletion (top) and inversion (bottom) variants (markers), stratified by occurrence in early or late time points (x-axis). Bars indicate mean and standard error of mean. (**B**) Gene essentiality scores (y-axis) across all genes and those associated with early and late inversions in the HAP1 cell line (x-axis). Bars indicate mean and standard error of mean.

**Supplementary Figure 13.**
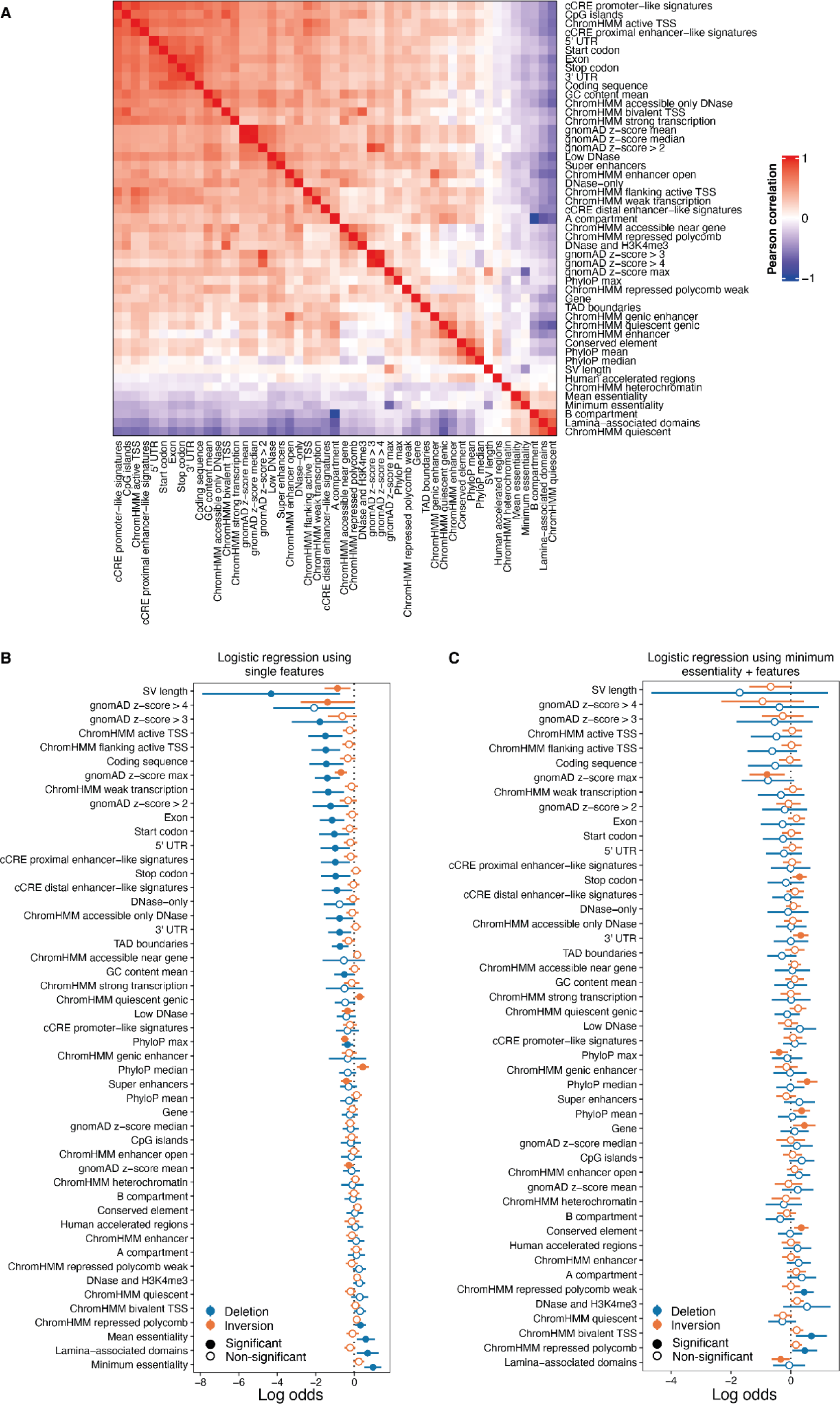
Features under selection in deletions and inversions. (**A**) Heatmap of correlations (colored by Pearson’s R) between various genomic and epigenomic features (x and y axes) and their frequency of occurrence within deletion and inversion structural variants. (**B**) Log odds (x-axis) for features (y-axis) between early variants and surviving variants colored by variant type (logistic regression). Markers represent the log odd and whiskers 95% confidence intervals. Unless indicated otherwise, the fraction of each variant covered by the respective feature is used as the statistic. (**C**) As (B) but using a logistic regression model that includes minimum essentiality as an additional variable.

**Supplementary Figure 14.**
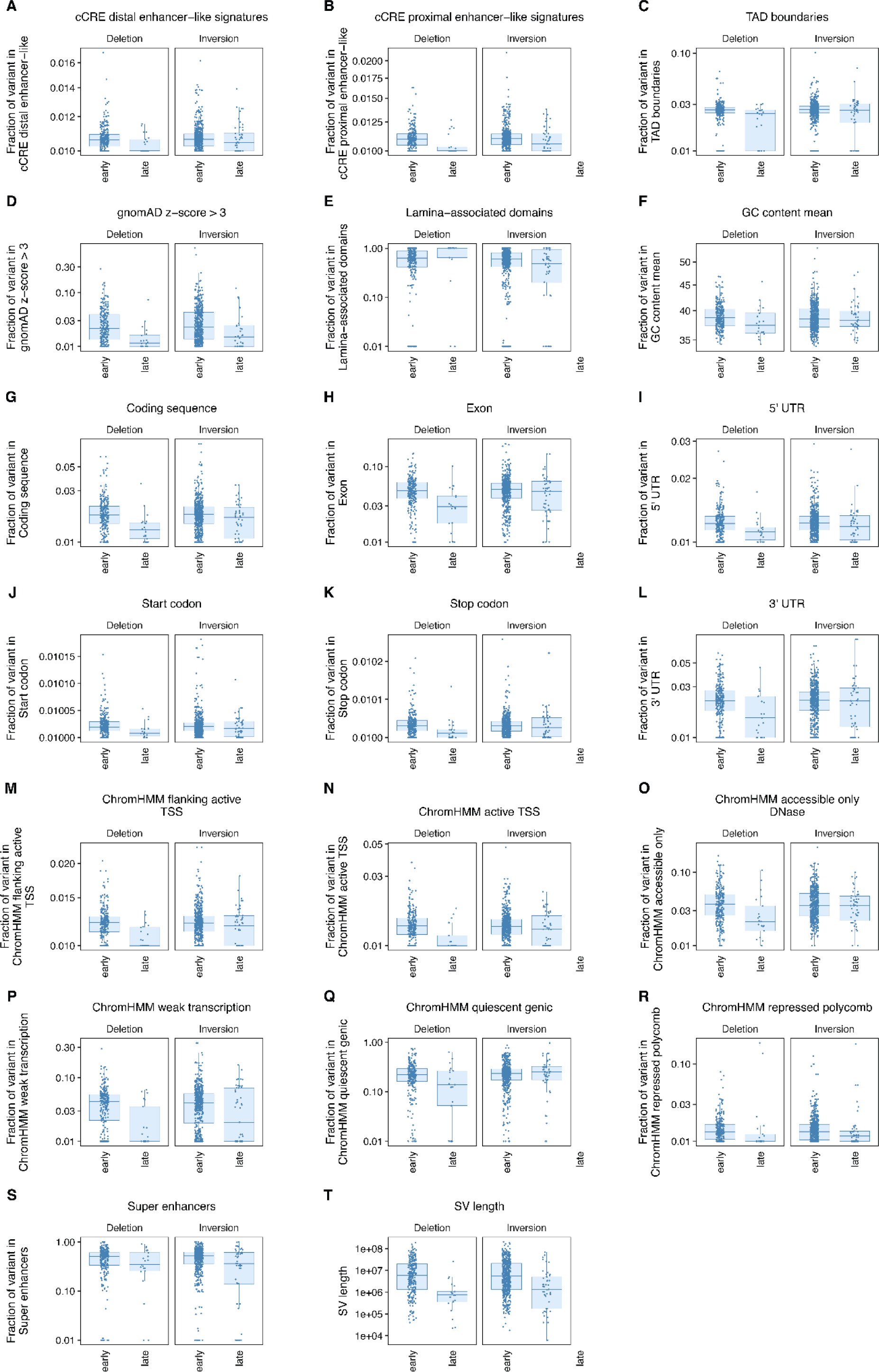
A variant level view on differences between early and late deletions and inversions. (**A-T**): Feature level (y-axis) for early and late time point deletions (x-axis) in HAP1 for each significant feature (from Figure 4d). Markers represent individual structural variants. Box: median and quartiles. Whiskers: Largest or smallest value no further than 1.5 interquartile ranges from the hinge.

**Supplementary Figure 15.**
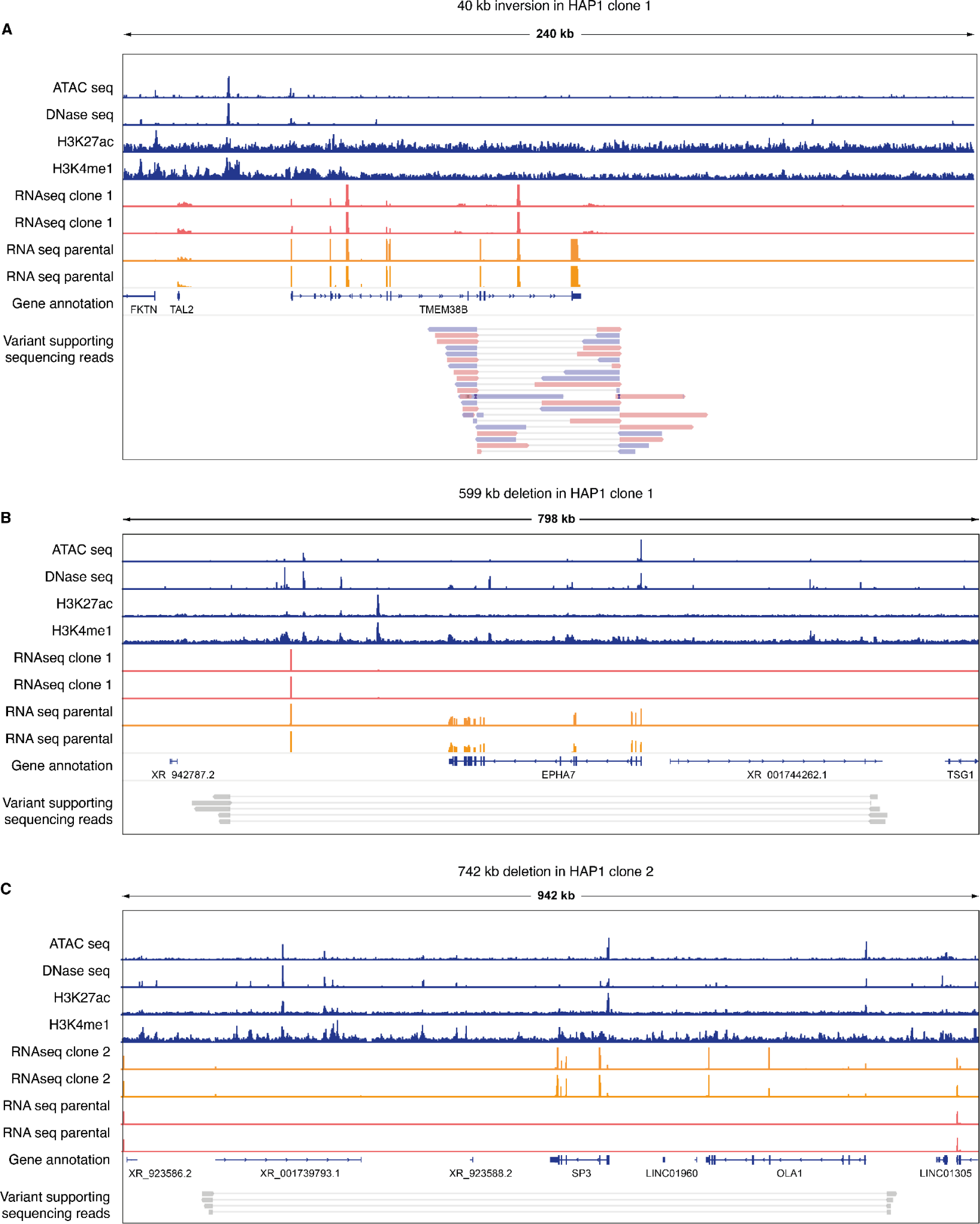
Genomic landscape of Cre-induced variants in HAP1 clones. (**A**) Integrated genomic profile displaying a 240 kb region encompassing a 40 kb inversion.Tracks from top to bottom represent ATAC-seq and DNase-seq data indicating open chromatin regions; histone modification marks H3K27ac and H3K4me1 associated with active regulatory elements; RNA-seq data from HAP1 clone 1 and parental lines; gene annotation; and variant supporting sequencing reads, with alignments depicting the inversion breakpoints. (**B**) As (A) but for a 798 kb region encompassing a 599 kb deletion in clone 1. (**C**) As (A) but for a 942 kb region encompassing a 742 kb deletion in clone 2.

**Supplementary Figure 16.**
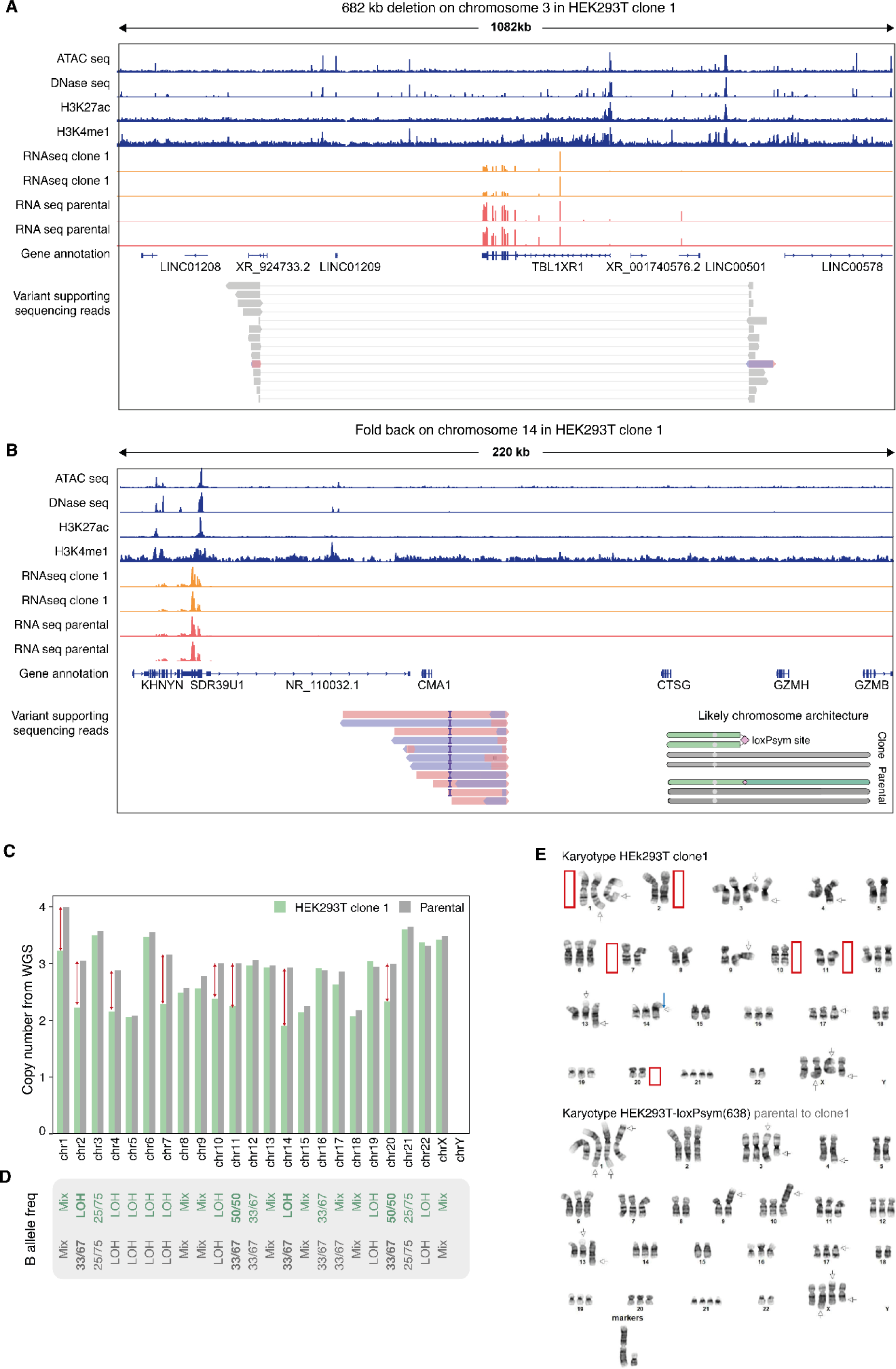
A HEK293T clone with an isochromosome. (**A**) Integrated genomic profile displaying a 1082 kb region encompassing a 692 kb deletion. Tracks from top to bottom represent ATAC-seq and DNase-seq data indicating open chromatin regions; histone modification marks H3K27ac and H3K4me1 associated with active regulatory elements; RNA-seq data from HEK293T clone 1 and parental lines; gene annotation; and variant supporting sequencing reads, with alignments depicting the inversion breakpoints. (**B**) As (A) but for a fold-back on chromosome 14. (**C**) Median copy number (y-axis, normalized to chromosomes 12 and 19 which are triploid in parental and clone) per chromosome (x-axis) for parental HEK293T-loxPsym(638) cells and a scrambled clone (colors). Red arrows indicate chromosomes with copy number differences > 0.5. (**D**) Most common B allele frequencies for the scrambled clone and parental per chromosome. ‘Mix’ for chromosomes where no single BAF made up at least 50% of the chromosome. (**E**) Representative G-band karyotype of parental and the scrambled clone (n = 23 metaphase spreads). Red boxes indicate chromosomes that are absent in the scrambled clone and arrows indicate translocations (the blue one points to the abnormal chromosome 14 in the scrambled clone).

